# Infection strategy and biogeography distinguish cosmopolitan groups of marine jumbo bacteriophages

**DOI:** 10.1101/2022.01.18.476781

**Authors:** Alaina R. Weinheimer, Frank O. Aylward

## Abstract

Recent research has underscored the immense diversity and key biogeochemical roles of large DNA viruses in the ocean. Although they are important constituents of marine ecosystems, it is often difficult to detect these viruses due to their large size and complex genomes. This is true for “jumbo” bacteriophages, which have genome sizes >200 kbp and large capsids reaching up to 0.45 μm in diameter. In this study, we sought to assess the genomic diversity and distribution of these bacteriophages in the ocean by developing a bioinformatic pipeline to generate and validate jumbo phage genomes from metagenomes. We recover 85 marine jumbo phages that ranged in size from 201-498 kilobases, and we examine their genetic similarities and biogeography together with a reference database of marine jumbo phage genomes. By analyzing Tara Oceans metagenomic data we show that jumbo phages are less common in <0.22 μm size fractions but are widespread in larger fractions, consistent with their large size. Our network-based analysis of gene sharing patterns reveals that jumbo bacteriophage belong to five genome clusters that are typified by diverse replication strategies, genomic repertoires, and potential host ranges. Our analysis of jumbo phage distributions in the ocean reveals that depth is a major factor shaping their biogeography, with some phage genome clusters exhibiting higher relative abundance in either surface or mesopelagic waters, respectively. Taken together, our findings demonstrate that jumbo phages are widespread community members in the ocean with complex genomic repertoires and ecological impacts that warrant further targeted investigation.

## Introduction

Although historically noted for their small virion sizes and simple genomes [1], viruses with large particles and elaborate genomes have been discovered in recent decades throughout the biosphere [2–4]. These complex viruses not only invite intriguing evolutionary questions [5–7], but also expand the potential roles viruses have in shaping microbial community structure and biogeochemical cycling [4, 8–10]. One group of these larger viruses are jumbo bacteriophages (jumbo phages), which have traditionally been defined as *Caudovirales* with genomes over 200 kilobases in length [11]. While a recent survey of cultured jumbo phages by Iyer and colleagues [6] showed jumbo phages share some universal features and genes, such as encoding DNA polymerases and the terminase large subunit (TerL), these features do not distinguish them from smaller phages, and several lines of evidence suggest that jumbo phages emerged from smaller phages multiple times independently. For example, a recent phylogenetic study of cultured *Caudovirales* used concatenated protein alignments to generate phylogenies and found that the most supported clades within the *Caudovirales* family do not consistently correspond to genome length [12]. Furthermore, Iyer et al. found that jumbo phages cluster with smaller phages based on gene content and are best grouped by replication machinery, among other infection apparati [6]. Taken together, jumbo phages likely emerged from diverse smaller phages multiple times independently and form distinct clades within the *Caudovirales*.

Although the first jumbo phages were isolated as early as the 1970s [13], these viruses have remained relatively sparse in culture, representing less than 3% (n=93) of complete *Caudovirales* genomes on NCBI’s RefSeq Viral Genome Portal (downloaded July 5, 2020). All cultured jumbo phage capsids have morphologies belonging to the *Myoviridae* and *Siphoviridae* families and undergo infection cycles that reflect temporal patterns of lytic *Caudovirales* [14]. Some jumbo phages are known to stall infections resulting in “pseudolysogeny”, however, which has been proposed as a competitive strategy against other phages to prevent superinfection [15]. Jumbo phages that have been studied extensively are primarily investigated for their unusually complex functional capabilities, such as encoding entire transcriptional apparatti [16] or sophisticated anti-CRISPR defense mechanisms [17, 18]. Although most cultivated jumbo phages have been isolated from a variety of Gram positive and Gram negative model and pathogenic bacteria [14], a recent metagenomic survey uncovered these viruses in diverse ecosystems around the world [4].

Despite this apparent broad environmental distribution, commonly-used methods for viral isolation and diversity surveys bias against the inclusion of jumbo phages. Because viruses have historically been considered to be smaller than cells, many viral diversity surveys specifically examine only small particle sizes. For example, in plaque assays, agar concentrations are often too high for larger phage particles to diffuse through compared to smaller particles [19]. Moreover, filters are often used to remove cells when preparing viral enrichments for metagenomic sequencing [20], which excludes larger viruses [9, 21]. Particularly in marine studies, the <0.22 μm fraction, sometimes even referred to as the “viral fraction” [22], is most commonly examined for viruses[23–25]. Jumbo phages can have particles over 0.45 μm in length (i.e. *Pseudomonas aeruginosa* phage PhiKZ)[13], however, and will therefore be excluded from <0.22 μm size fractions. Lastly, in bioinformatic pipelines, phage sequences are typically only assembled to the contig or scaffold level, which is sometimes sufficient for the assembly of most known smaller phage genomes [26], but often leaves larger phage genomes fragmented into multiple contigs and may require additional joining of contigs into bins [27]. Overall, considering these biases and the recently-discovered broad distribution of these viruses [4], jumbo phages may represent underappreciated components of marine microbial communities and food webs that warrant further examination.

In this study, we examine the diversity and prevalence of jumbo phages in the global ocean. We developed a workflow for generating and validating high-quality jumbo bacteriophage bins from metagenomic data, with which we identified 85 bins of jumbo phages. We then compared the genetic content of these jumbo phages with other cultured phages of all sizes and metagenomic jumbo phages from other studies. We found that the jumbo phages of this study grouped into five distinct clusters that are distinguished by diverse replication machinery and infection strategies, implicating a broad range of potential jumbo-phage host interactions in the ocean. We then assessed the distribution of jumbo phages belonging to these genome clusters in the ocean by using metagenomic data from Tara Oceans [28,29]. Mapping the Tara Oceans metagenomic data onto the jumbo phage sequences revealed that these jumbos phages are collectively widely distributed in the ocean, but vary in biogeography based on cluster, with some more enriched in surface waters relative to deeper waters and vice versa. Upon examining the collective presence of jumbo phages in different filter fractions, we also found that commonly used size fractions in viral diversity surveys reduce the ability to recover and detect jumbo phage sequences. As these complications apply to many viral diversity studies in other environments, our results indicate that jumbo phages may play underappreciated roles in ecosystems around the globe.

## RESULTS AND DISCUSSION

### Detection and validation of high-quality jumbo phage bins

Due to the large size of jumbo bacteriophage genomes, it is likely that they are present in multiple distinct contigs in metagenomic datasets and therefore require binning to recover high-quality metagenome-assembled genomes (MAGs) [27]. This has been shown for large DNA viruses that infect eukaryotes, where several recent studies have successfully employed binning approaches to recover viral MAGs [2, 3]. Here, we used the same 1,545 high-quality metagenomic assemblies [30] used in a recent study to recover giant viruses of eukaryotes [3], but we modified the bioinformatic pipeline to identify bins of jumbo bacteriophages. We first binned the contigs from these assemblies with MetaBat2 [31], which groups contigs together based on similar nucleotide composition and coverage profiles, and focused on bins of at least 200 kilobases in total length. We subsequently identified bins composed of bacteriophage contigs through analysis with VirSorter2 [32], VIBRANT [33], and CheckV [34] (see Methods for details).

The occurrence of multiple copies of highly-conserved marker genes is typically used to assess the level of contamination present in metagenome-derived genomes of bacteria and archaea [35]. Because bacteriophage lack these marker genes, we developed alternative strategies to assess possible contamination in our jumbo phage bins. Firstly, we refined the set of bins by retaining only those with no more than 5 contigs that were each at least 5 kilobases in length to reduce the possibility that spurious contigs were put together. Secondly, we assessed the possibility that two strains of smaller phages with similar nucleotide composition may be binned together by aligning the contigs in a bin to each other. Bins that had contigs with high sequence similarity across the majority of their lengths were discarded (Supplementary Figure 1). Thirdly, we discarded bins if their contigs exhibited aberrant co-abundance profiles compared to other contigs in the same bin. We identified these bins through read mapping and co-abundance analysis of 225 marine metagenomes from Tara Oceans. Coverage variation was benchmarked based on read-mapping results from artificially-fragmented reference genomes present in the samples (See Methods for details). Only bins with coverage variation below our empirically-derived threshold were retained. Using this stringent filtering, we identified 85 bins belonging to jumbo bacteriophages that are present in Tara Oceans metagenomes. These bins ranged in length from 202 bp to 498 kbp, and 31 consisted of a single contig, while 54 consisted of 2-5 contigs (Supplemental Figure 2).

To assess global diversity patterns of jumbo bacteriophage we combined these jumbo phage bins together with a compiled database of previously-identified jumbo phages that included all complete jumbo *Caudovirales* genomes on RefSeq (downloaded July 5th, 2020), the INPHARED database [36], the Iyer et al 2021 study [6], the Al-Shayeb et al. 2020 study [4], and marine jumbo phage contigs from metagenomic surveys of GOV 2.0 [25] (60 jumbo phages), ALOHA 2.0 [37] (8 jumbo phages), and one megaphage MAG recovered from datasets of the English Channel [38]. Ultimately, we arrived at a set of 223 jumbo phages present in at least one Tara Oceans sample that we analyzed further in this study and refer to as marine jumbo phages. Statistics on genomic features can be found in Supplemental Dataset 1.

### Marine jumbo phages belong to distinct groups with diverse infection strategies

Because bacteriophages lack high-resolution, universal marker genes for classification, such as 16S rRNA in bacteria, phages are often grouped by gene content [39, 40]. Here, we generated a bipartite network that included the 85 bins of jumbo phages with a dataset of available *Caudovirales* complete genomes in RefSeq (3,012 genomes; downloaded July 5th, 2020) and the set of jumbo phages described above. To construct the bipartite network, we compared proteins encoded in all the phage genomes to the VOG database (vogdb.org), and each genome was linked to VOG hits that were present (Figure 1, Supplemental Dataset 2, see Methods for details). To identify groups of phage genomes with similar VOG profiles, we employed a spinglass community detection algorithm [41] to generate genome clusters. The marine jumbo phages of this study clustered into five groups that all included both jumbo and non-jumbo phage genomes (Figure 2a). We refer to these five clusters as Phage Genome Clusters (PGCs): PGC_A, PGC_B, PGC_C, PGC_D, and PGC_E. These PGCs included cultured phages and metagenome-derived jumbo phages found in various environments (i.e. aquatic, engineered) and isolated on a diversity of hosts (i.e. Firmicutes, Proteobacteria, Bacteroidetes) (Figure 2b,c). Of the marine jumbo phages, 134 belonged to PGC_A, 11 to PGC_B, 90 to PGC_C 27, 7 to PGC_D, and 1 to PGC_E (Figure 1b). In addition to this network based analysis, we also examined phylogenies of the major capsid protein (MCP) and the terminase large subunit (TerL) encoded by the marine jumbo phages and the same reference phage set examined in the network (Figure 1c, 1d). With the exception of PGC_A, the marine jumbo phages that belong to the same PGC appeared more closely related to each other than those belonging to different clusters. The evolutionary history of these marker genes therefore supports the results of the bipartite network genome clustering and is consistent with the view that genome gigantism evolved multiple times independently within the *Caudovirales* [6].

**Figure 1.**
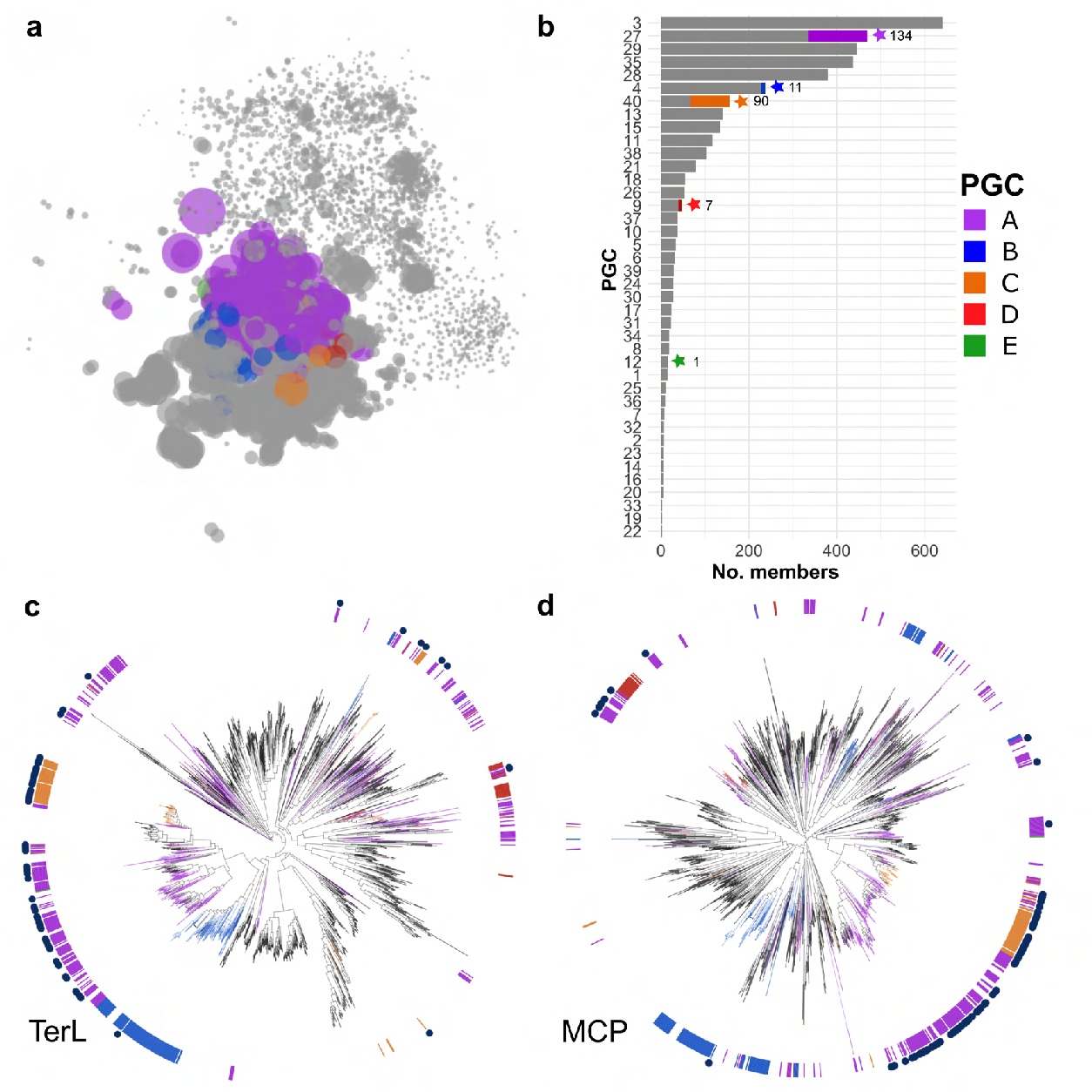
Comparison of bins to reference gene content and evolutionary histories. **(a)** Network with marine jumbos and references as nodes and edges based on shared VOGs. Marine jumbo phage nodes are colored by PGC as detected with spinglass community detection analysis, other nodes are in gray. (**b)** Barchart of the number of members in each PGC. PGCs with marine jumbo phages are denoted with a star and the number of marine jumbo phages in that PGC. Proportion of marine jumbo phages in that PGC is colored. (**c,d)** Phylogenies of TerL (**c**) and MCP (**d**) proteins with references and bins. Inner ring and branches are colored by the 5 PGCs that marine jumbo phages belong to. Navy blue circles in the outer ring denote marine jumbo phages.

**Figure 2.**
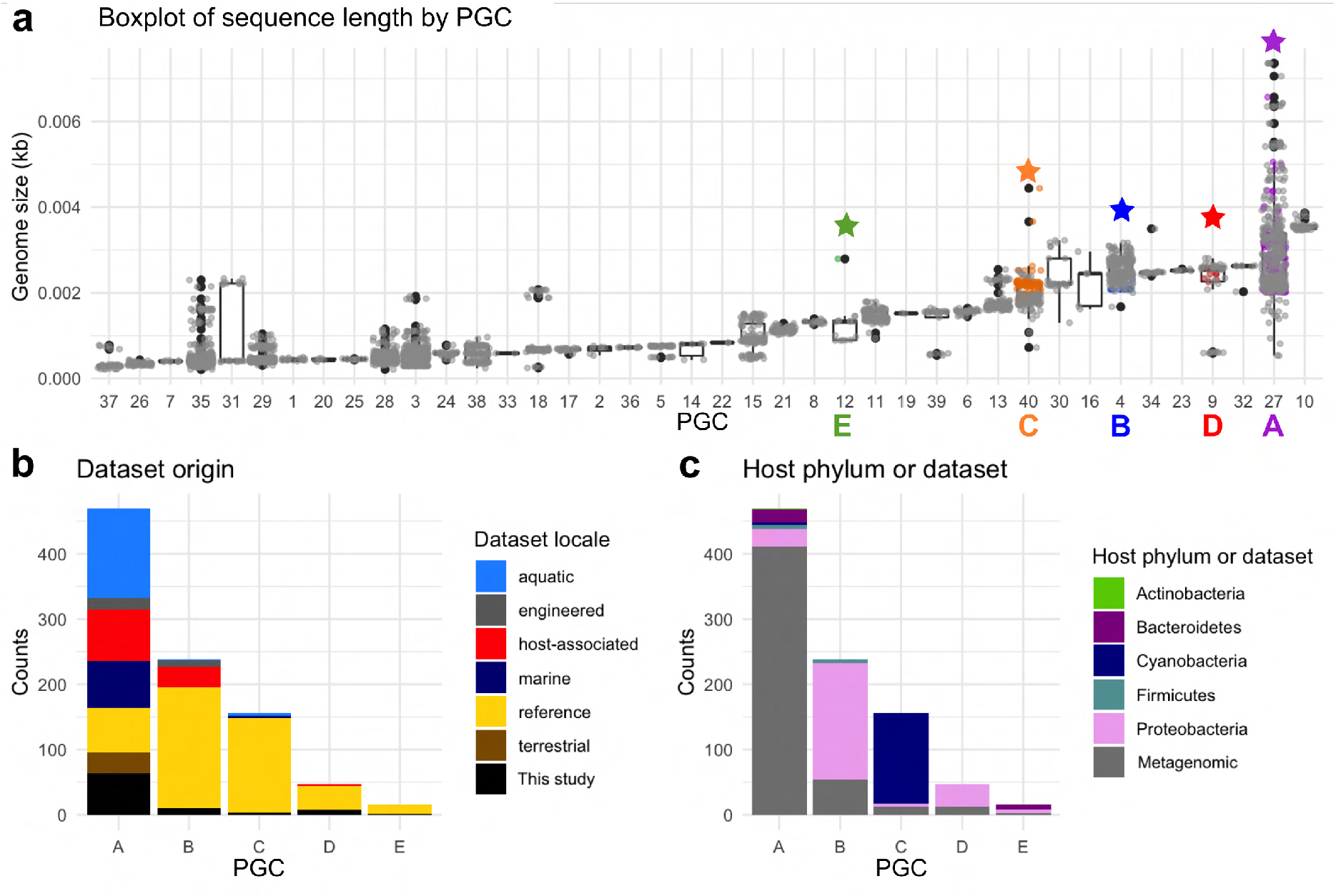
**(a)** Boxplot of genome length in each network cluster (x-axis is PGC number). Star denotes PGC with a marine jumbo phage and the color matches the PGC letters of Figure 1. **(b)** Stacked barplot of the metagenome environment from which each phage derives from in each PGC (x-axis). Reference (yellow) are cultured phages, black is the bins of jumbo phages from this study **(c)** Stacked barplot of the host phylum of the RefSeq cultured phages in each cluster; metagenomic phages are in gray.

We then compared functional content encoded by the marine jumbo phages in the PGCs to identify functional differences that distinguish these groups. PGC_E was excluded from this analysis because this genome cluster contained only a single jumbo phage. Collectively, most genes of the marine jumbo phages could not be assigned a function (mean: 86.60%, std dev: 7.01%; Supplemental Dataset 3), which is common with environmental viruses [42, 43]. Genes with known functions primarily belonged to functional categories related to viral replication machinery, such as information processing and virion structure (Figure 3a), and these genes drove the variation between the genome clusters of marine jumbo phages (Figure 3b). Iyer et al provided a comparative genomic analysis that was able to identify three types of jumbo phages that are defined by different infection strategies and host interactions (referred to as Groups 1-3) [6]. We cross-referenced our PGCs and found that PGCs B, C, and D of this study corresponded to Iyer Groups 1, 2, and 3, respectively, suggesting that these genome clusters contain phages with distinct infection and replication strategies. PGC_A corresponded to multiple phage groups in the Iyer study, indicating that this genome cluster contains a particularly broad diversity of phages.

**Figure 3.**
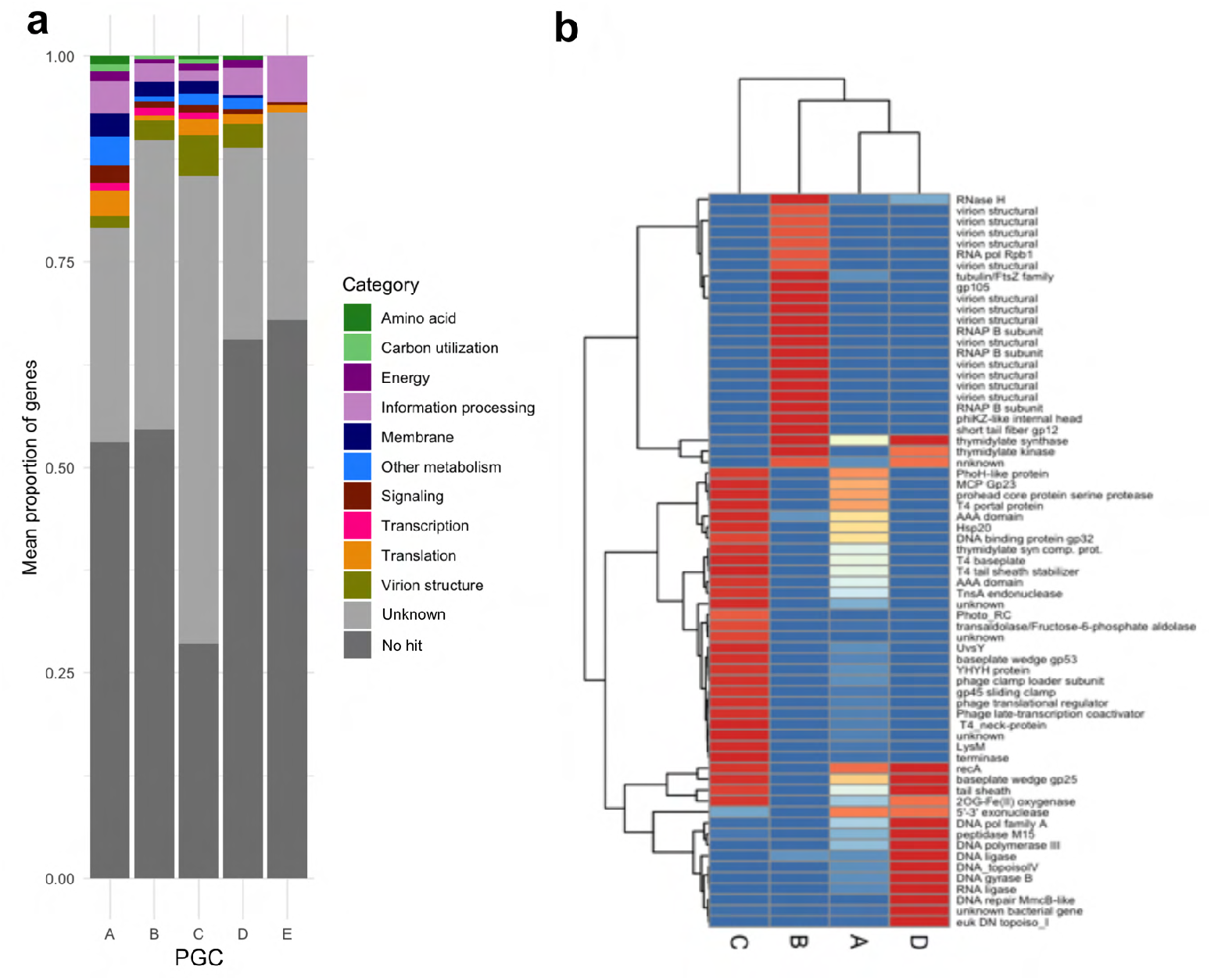
(**a**) Functional categories for genes encoded by jumbo phages. (**b**) Heatmap of proportion of genomes in each cluster that contain the listed genes. Listed genes were selected based on containing a known function and variance between the clusters above 0.2. Dendrogram was generated based on hierarchical clustering in pheatmap.

PGC_B consists of 238 phages (11 (4.6%) marine jumbo phages, including 10 bins generated here), and included cultured phage of Group 1 of Iyer et al, which is typified by *Pseudomonas aeruginosa* phage PhiKZ. Supporting this correspondence with Iyer Group 1, all marine jumbo phages of PGC_B encoded the same distinct replication and transcription machinery, including a divergent family B DNA polymerase and a multi-subunit RNA polymerase (Figure 3b, Supplemental Dataset 3). These marine jumbo phages also encoded a PhiKZ internal head protein, and they uniquely encoded shell and tubulin homologs which has recently been found in PhiKZ phages to assist in the formation of a nucleus-like compartment during infection that protects the replicating phage from host defenses [17,18]. Although we could not confidently predict hosts for the 11 metagenomic marine jumbo phages in this PGC_B (Supplemental Dataset 1), the cultured phages of this genome cluster infect pathogenic bacteria belonging to the phyla Proteobacteria (178 phages) and Firmicutes (6 phages) (Figure 2c), implicating a potential host range for marine jumbo phages in PGC_B.

PGC_C is comprised of 156 phages total (90 marine jumbo phages (57.7%); 4 bins generated from this study) and included reference jumbo phages in Iyer Group 2 (31, 19.9%) which are typified by Alphaproteobacteria and Cyanobacteria phages. Likewise, the host range of other cultured phages in PGC_C support the Group 2 correspondence, either infecting Cyanobacteria (139 phages) or Proteobacteria (4 phages) (Figure 2c). Furthermore, all 3 marine metagenomic phages in PGC_C for which hosts could be predicted were matched to Cyanobacteria hosts (Supplemental Dataset 1). Functional annotations of PGC_C marine jumbo phages revealed nearly all encode a family B DNA polymerase (97.8% phages) and the photosystem II D2 protein (PF00124, VOG04549) characteristic of cyanophages (90% phages) (Figure 3b). This PGC included the reference *Prochlorococcus* phage P-TIM68 (NC_028955.1), which encodes components of both photosystem I and II as a mechanism to enhance cyclic electron flow during infection [44]. This suggests that an enhanced complement of genes used to manipulate host physiology during infection may be a driver of large genome sizes in this group. Additionally, most of the PGC_C marine jumbo phages encoded Lipopolysaccharide (LPS) biosynthesis proteins (76%), which have been found in cyanophage genomes that may induce a “pseudolysogeny” state, when infected host cells are dormant, by changing the surface of the host cell and preventing additional phage infections [15] (Supplemental Dataset 3). Taken together, most marine jumbo phages of PGC_C likely follow host interactions of jumbo cyanophages, such as potentially manipulating host metabolism by encoding their own photosynthetic genes and potentially inducing a pseudolysogenic state.

PGC_D comprised 47 phages, of which 7 were marine jumbo phages generated in this study (14.9%). This group included Iyer Group 3 jumbo phages (15, 31.9%), which is primarily distinguished by encoding a T7-type DNA polymerase but is not typified by a particular phage type or host range. Supporting this grouping, all marine jumbo phages in this study encoded this T7 DNA polymerase which belongs to family A DNA polymerases (Figure 3b, Supplemental Dataset 3). Most of the other genes distinctively encoded by the marine jumbo phages in this group included structural genes related to T7 (T7 baseplate, T7 capsid proteins), a eukaryotic DNA topoisomerase I catalytic core (PF01028), and a DNA structural modification genes (MmcB-like DNA repair protein, DNA gyrase B). Hosts of metagenomic marine jumbo phages in PGC_D could not be predicted (Supplemental Dataset 1); however, cultured Group 3 jumbo phages in PGC_D all infect Proteobacteria, primarily Enterobacteria and other pathogens.

PGC_A was the largest of the PGCs with 469 phages in total, including 134 marine jumbo phages (63 bins from this study). This genome cluster contained the largest jumbo phages, such as *Bacillus* phage G (498 kb) and the marine megaphage Mar_Mega_1 (656 kb) recently recovered from the English Channel [38]. Unlike other PGCs, PGC_A contained mostly metagenomic phages (401, 85%, Figure 2b,c). Considering PGC_A contains the largest jumbo phages (Figure 1b, 2a), the vast genetic diversity in this PGC might explain why few genes were found to distinguish this group. Of the genes unique to PGC_A, only one was present in at least half of the phages (51.9%), which was a Bacterial DNA polymerase III alpha NTPase domain (PF07733). The host ranges of cultured phages from this PGC further reflect the phage diversity and included a variety of phyla and genera that can perform complex metabolisms or lifestyles, such as the nitrogen-fixing Cyanobacteria of the *Nodularia* genus isolated from the Baltic Sea (corresponding to phage sequences NC_048756.1 and NC_048757.1) and the Bacteroidetes bacteria *Rhodothermus* isolated from a hot spring in Iceland [45].

### Relative abundance of jumbo bacteriophages across size fractions

To explore the distribution of the marine jumbo phages in the ocean, we first examined the size fractions in which the jumbo phages were most prevalent. To remove redundancy, we clustered the 223 jumbo phages into populations if they were highly similar at the nucleotide level ( >80% genes with >95% average nucleotide identity) [23], which resulted in 120 jumbo phage populations. Out of the 225 Tara Oceans metagenomes examined, 202 (89.8%) contained at least one jumbo phage population (median: 24, Supplemental Dataset 4). Jumbo phages were less frequently detected in samples below 0.22 μm (<-0.22 μm, 0.1-0.22 μm) than those above 0.22 μm (0.45-0.8 μm, 0.22-0.45 μm, 0.22-1.6 μm, 0.22-3 μm). Only 54.5% of samples in the <-0.22 μm fraction and 82.4% of samples in the 0.1-0.22 μm fraction had at least one jumbo phage present. Conversely, all samples in all larger size fractions above 0.22 μm had at least one jumbo phage present (Figure 4a). Furthermore, we detected 21 populations (14.9%) exclusively in samples above 0.22 um, compared to only one population (0.71%) exclusive to samples below 0.22 μm. A similar disparity in virus detection between size fractions has been reported for large eukaryotic viruses, where roughly 41% of phylotypes were present in samples of the picoplankton fraction of 0.22-3 μm but absent in size fractions below 0.22 μm [9]. In contrast to this study, where certain viral groups were more prevalent in larger size fractions than smaller, a jumbo phage’s PGC membership or genome size did not affect its probability of detection at different size fractions (Supplemental Figure 3, Supplemental Figure 4).

**Figure 4.**
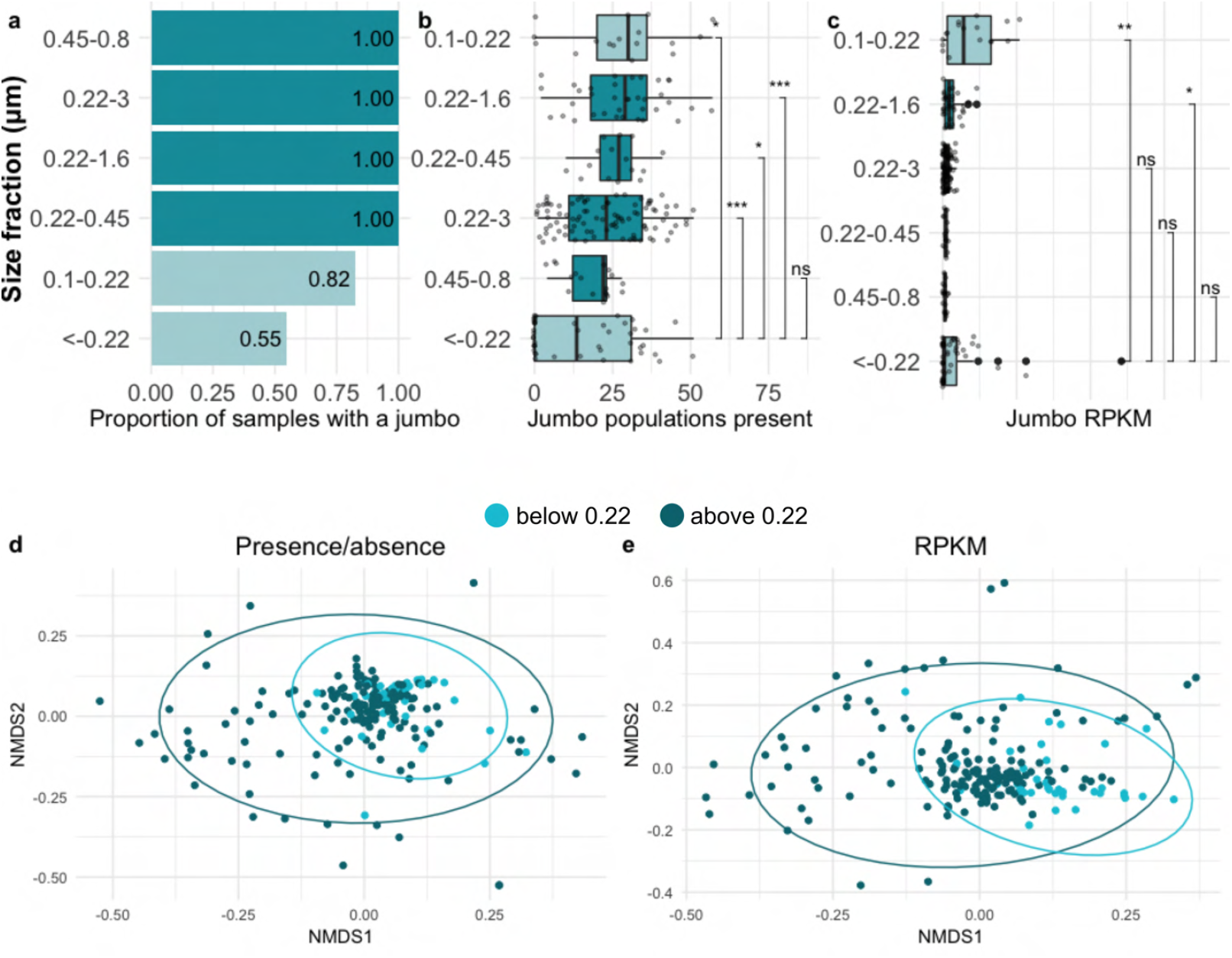
Comparison of jumbo abundance and presence in samples of different filter size fractions. Dark teal are fractions with minimum sizes of 0.22 μm or higher. Light teal are fractions with a maximum size of 0.22 μm or lower. **a)** Barchart of the proportion of samples with at least one marine jumbo phage (x-axis) by size fraction (y-axis) sorted from highest to lowest. **b)** Boxplot with x-axis as the number of marine jumbo phages found in a sample with size fraction on the y-axis sorted by median. **c)** Boxplot with x-axis as the relative abundance of marine jumbo phages found in a sample (RPKM) with size fraction on the y-axis sorted by median. Significance bars in c,d correspond to Wilcox test, with stars corresponding to p-values < 0.05 and those with p-values > 0.05 as not significant “ns” (stat_compare_means function). (**d,e**) NMDS plots (Bray Curtis dissimilarity distances) of jumbo phage composition in each sample using presence absence data (**d**) and relative abundance data (**e**). Samples are colored by size fraction distinction above 0.22 (dark teal) and below 0.22 (light teal). Ellipses calculated based on multivariate normal distribution. Samples above and below 0.22 were not significantly different from each other using ANOSIM (p-values > 0.05).

We also compared jumbo phage diversity (defined as population richness), relative abundance, and community composition between the size fractions. Samples of the size fraction <-0.22 μm were significantly less diverse (P-value 0.0003, Wilcox test) and had significantly lower relative abundances of jumbo phages relative to the large size fractions of 0.1-0.22 and 0.22-1.6 (P-values 0.003, 0.031 Wilcox test, respectively). Samples of the 0.1-0.22 μm size fraction had the highest diversity and relative abundances compared to the other fractions (Figure 4b,c). The enhanced signal of jumbo phages relative to other entities in the 0.1-0.22 μm size fraction may result from the reduction of cells below the 0.22 μm threshold and the reduction of most other viruses above the 0.1 μm limit. Despite these differences in diversity and relative abundances, jumbo phage community composition did not significantly differ between the >0.22 and <0.22 μm size fractions (P-value 0.8109, ANOSIM, presence/absence Bray-Curtis distance matrix, Figure 4d,e).

To directly test the effect of the 0.22 μm size fraction cut-off on jumbo phage recovery, we examined a subset of the samples that were co-collected at the same station and depth for the fractions below 0.22 μm (<-0.22 or 0.1-022) and above 0.22 μm (0.1-0.22 μm or 0.22-3 μm). The number of detected jumbo phage populations was significantly higher in samples above 0.22 μm than below 0.22 μm (P-value = 0.0004, Wilcox test, Supplemental Figure 5a). The relative abundance of jumbo phages was not significantly different between this size distinction (P-value = 0.1309, Wilcox test, Supplemental Figure 5b), however, nor was community composition (P-value = 0.1229, ANOSIM, presence/absence Bray-Curtis distance matrix, Supplemental Figure 5c,d). Taken together, these findings suggest that using size fractions below 0.22 μm to analyze phages limits the recovery of jumbo phage populations detected, likely due the large size of these viruses. The enhanced relative abundance of jumbo phages in the 0.1-0.22 μm relative to the other size fractions suggests that this may be the optimal size fraction to examine for studies that seek to target jumbo phages specifically.

### Biogeography of jumbo bacteriophages in the global ocean

#### Jumbo phage populations varied in depth distribution by PGC

Jumbo populations in this study were found in all three Longhurst biomes (Trades, Westerlies, Coastal) and depths (Surface, Deep Chlorophyll Maximum, Mesopelagic) examined (Figure 5, Supplemental Figure 6; Supplemental Dataset 4). We focused these biogeographic analyses on the 0.22-1.6 or 0.22-3 μm size fractions because the most sites were available for these samples, and because we found that jumbo phages were recovered at a higher frequency in larger size fractions. Jumbo phage communities differed significantly between depths (P-value = 0.0001, ANOSIM based on presence/absence Bray-Curtis distance matrix, Supplemental Figure 7c,d), consistent with the dramatic transition in community composition that occurs from surface waters to below the deep chlorophyll maximum [37, 46]. Specifically, the diversity of jumbo phages across depths varied by genome cluster, with PGC_A and PGC_C exhibiting higher abundance and diversity in the epipelagic (SRF and DCM), and PGC_B and PGC_D exhibiting higher abundance in the mesopelagic (Figure 5d, Supplemental Figure 7a,b). PGC_C is typified by cyanophages, providing a clear reason why this phage group is enriched in surface waters. Conversely, PGC_B is typified by *Pseudomonas aeruginosa* PhiKZ phages, suggesting these PGC_B marine jumbo phages may be infecting heterotrophic bacteria that potentially thrive in deeper waters. Overall relative abundance and diversity of jumbo phages in this study were significantly higher in the epipelagic zone, partly because most of these viruses are in PGC_A. This pattern held when examining only those samples that were co-collected at all three depths (Supplemental Figure 8,9). This general pattern therefore reflects what has been found in previous studies on the depth distribution and viral protein clusters, where more were unique in the euphotic (i.e. epipelagic) than aphotic depths [9, 25, 47, 48], although this contrasts what has recently been found in the Pacific Ocean, where overall viral diversity increased in the mesopelagic [37].

**Figure 5.**
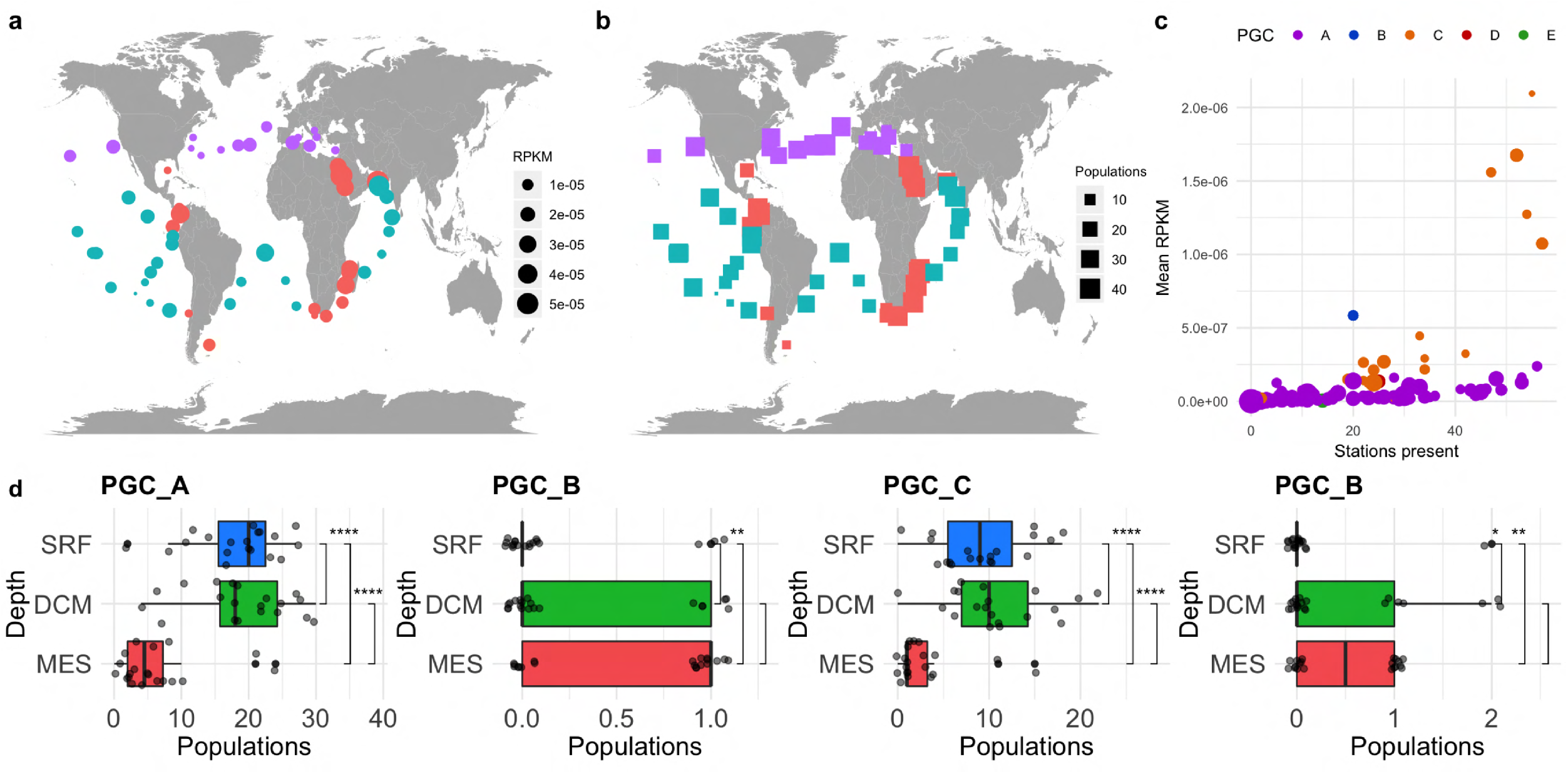
**a,b**) Maps of the relative abundance (**a**) of jumbo phages (in RPKM) and (**b**) number of jumbo populations present in each surface (SRF) sample of the picoplankton size fraction (either 0.22-3μm or 0.22-1.6μm depending on availability). Dots sizes are proportional to the number of populations or RPKM and colored by biome (Coastal - pink, Westerlies - purple, Trades - blue). (**c**) Scatterplot of the mean RPKM of a jumbo population in SRF picoplankton samples versus the number of SRF picoplankton stations it was present. Populations are colored by PGC and size corresponds to putative genome length in 100 kilobases. (**d**) Boxplot of the number of jumbo phage populations in a sample separated by depth sorted by median for each PGC. Significance bars correspond to Wilcox test, with stars corresponding to p-values < 0.05 and those with pvalues > 0.05 as not significant (stat_compare_means function)

#### Jumbo phage biogeography across biomes

Jumbo phage communities in this study significantly differed in composition in surface samples between the Trades, Coastal, and Westerlies biomes, albeit weakly (P-value 0.0021, ANOSIM, presence/absence Bray-Curtis distance matrix, Supplemental Figure 10c,d). However, no biome appeared to be a particular hotspot for jumbo phages, as they were not significantly more diverse in any particular biome (Supplemental Figure 10a). Although the relative abundances of jumbo phages were higher in Trades samples relative to Coastal samples (Supplemental Figure 10b), this pattern did not always hold when comparing biomes at different depths (Supplemental Figure 11). Similarly, jumbo phages from all PGCs could be detected in all biomes, but no PGC was consistently enriched in diversity and relative abundance in a particular biome (Supplemental Figure 12). A recent global study on marine viruses has found that viral diversity was better explained by ecological zones defined by physicochemical factors like temperature, rather than by Longhurst biomes defined by patterns of chlorophyll *a* concentrations [25], suggesting that Longhurst biomes may not be good predictors of viral diversity in general.

Jumbo phage populations ranged in endemicity with seven populations (5.8%) detected in only one Tara station and eleven (9.2%) present in 50 or more stations (Figure 5c). More endemic populations, present in only one Tara station, belonged to PGCs A, B, and C, while the most prevalent jumbo phage, those present in 50 or more stations, belonged to PGC_A and PGC_C. PGC_A and PGC_C contained the most populations, which likely explains the wide range of endemicity of phages in these clusters. Moreover, the cyanobacterial hosts that are known for many of the jumbo phages in PGC_C are widespread in the ocean, which may also explain the prevalence of this group of phages. In general, the heterogeneous distribution and abundance of these jumbo phages is consistent with the seed bank hypothesis, which postulates that viruses are passively dispersed throughout the ocean and viral community structure is shaped by local selective forces [23, 49]. This framework has previously been used to explain why phage distributions range from extremely cosmopolitan to extremely rare, which is a pattern that also appears to hold for jumbo bacteriophages.

## CONCLUSION

Large DNA viruses are becoming increasingly recognized as critical components of the virosphere, notable for their intriguing evolutionary histories [50, 51], vast functional capacities [3, 4], and global distribution [2, 4]. Here, we uncover the diversity and ecology of marine jumbo bacteriophages, which have historically been difficult to study due to biases in filtration and isolation strategies commonly used in virology. We employed a binning strategy to generate and quality-check genomes of jumbo phages and used it to identify 85 high quality bins. We combined these bins together with reference jumbo bacteriophage genomes, and ultimately identified 120 populations that are present in Tara Oceans metagenomes. When compared with other metagenomic jumbo phages and cultured phages of all sizes, we found that marine jumbo phages primarily belong to five phage genome clusters (PGCs) that encode distinct replication machinery, biogeography, and potential hosts. For example, marine jumbo phages in PGC_C follow cyanophage infection strategies and ecology, as this cluster included cultured marine cyanophages and encoded classic family B DNA polymerases and photosynthesis enzymes characteristic of cyanophages. Furthermore, we found they are enriched in surface waters relative to the mesopelagic, consistent with the geographic range of their hosts. In contrast, marine jumbo phages of PGC_B included cultured PhiKZ phages of *Pseudomonas aeruginosa* and uniquely encoded multi-subunit RNA polymerases and tubulin, which are thought to play a role in the remarkable nucleus-like structures that these viruses employ as an anti-CRIPSR defense [17, 18]. PGC_B was particularly enriched in mesopelagic waters, suggesting that this complex infection strategy may be more common in the deep ocean. These results demonstrate the diverse biology and ecology of jumbo phages, consistent with the hypothesis that jumbo phages evolved from smaller phages on independent occasions [6].

We show that jumbo phages are widespread throughout the ocean and are typically more diverse and abundant in epipelagic waters, which reflect previous findings that surface waters usually harbor a higher per-sample alpha diversity of viral groups compared to deeper waters [9, 25]. Larger phages therefore appear to coexist in patterns broadly similar to smaller viruses despite the disadvantages of their size, such as smaller burst sizes and lower host contact rates [52]. In eukaryotic giant viruses it has been hypothesized that these disadvantages are offset by higher infection efficiency, broader host ranges, potentially decreased decay rates, and higher rates of successful attachments compared to smaller viruses [52]. Although some of these advantages to viral gigantism may also apply to jumbo bacteriophages, it is unlikely that they are all applicable. For example, given that they are tailed *Caudovirales* [6], they likely possess higher host specificity in part due to their non-phagocytotic mode of infection. Nonetheless, the large genomes of jumbo bacteriophage often encode an expanded complement of genes used to manipulate host physiology during infection, and these may play critical roles in promoting infection efficiency or offsetting host defense mechanisms. The impressive complement of photosynthesis genes in PGC_C is at least partially responsible for the large genomes in this lineage, while the genes involved in anti-CRISPR defense found in PGC_B indicate that a host-virus arms race may be responsible for genome gigantism in this group. Interestingly, the largest number of jumbo phage genomes we identified belong to PGC_A, which is largely uncharacterized and composed of primarily metagenome-derived genomes, suggesting that these viruses have as-yet unidentified infection strategies. Overall, it is likely that the factors leading to genome gigantism in each of these genome clusters are different, given that they represent distinct lineages of *Caudovirales* that have independently evolved large genomes. Future work further characterizing the hosts of these jumbo phages and the details of their infection program, particularly in PGC_A, will therefore be critical to understanding mechanisms that underlie complexity in the virosphere and maintain diversity.

## Methods Summary

### Jumbo phage bin detection

Metagenomic scaffolds were downloaded from 1,545 assembled by Parks et al. 2017 [30] and binned with MetaBAT2 [31] (-s 200000 --unbinned -t 32 -m 5000 – minS 75 --maxEdges 75). Bins were retained if they summed to at least 200,000 base-pairs and comprised <= 5 contigs (min. contig size 5kb). Proteins were predicted with prodigal [53] using default settings on each bin individually. Bins were retained if they lacked more than one ribosomal protein, lacked overlapping regions (via promer and gnuplot [54] with MUMmer 3.0 [55]), had fewer hits to NCLDV than phage (via LASTp [56] against RefSeq r99), lacked more than one NCLDV marker gene (via HMM search (hmmer.org) against NCLDV marker gene HMM profiles [57]) and had even read coverage of Tara Ocean metagenomes via coverM [58] (https://github.com/wwood/CoverM). Jumbo phages were then detected by running the bins through VirSorter2 [32], VIBRANT [33], and CheckV [34]. Bins were considered putative phages if they had at least an average dsDNAphage score of >0.9 from VirSorter2 or VirSorter2 average score >0.5 and either i) CheckV quality of medium or higher (via pseudocontigs) or ii) VIBRANT consensus classification as viral. 85 bins were retained for downstream analyses. Bins were clustered into populations with a compiled set of jumbo bacteriophages (RefSeq phages over 200 kilobases, phage sequences over 200 kilobases from the INPHARED database [36], Iyer et al 2021 [6], Al-Shayeb et al. 2020 [4] jumbo phage genomes, GOV 2.0 [25], ALOHA 2.0 [37], and a megaphage assembled from the English Channel [38]) based on nucleotide sequences of genes (predicted with prodigal -d flag) aligned with BLASTn [59] (>95% average nucleotide identity, >80% genes) [23]. See Supplemental Methods for details.

### Evolutionary analyses

Reference phage sequences were compiled from RefSeq’s *Caudovirales* complete genomes (downloaded July 2020 from NCBI’s Virus genome portal; 3,012 genomes) along with the curated jumbo phage set used in the population analysis. Proteins of jumbo bins and this reference set were predicted with prodigal and searched against the Virus Orthologous Groups (VOGs, vogdb.org) via HMM searches (E-value 0.001). Bipartite network was made based on shared VOGs using igraph (graph.incidence) (1.2.5) [60] in R (version 3.5.1) [61] with RStudio (version 1.1.456) [62]. Clusters were detected by the spinglass community detection algorithm [41] via igraph (50 spins for 100 iterations). Final clusters determined by iteration with highest modularity. Phages were further classified with vContact2 [39]; however, none of the jumbo phage bins belonged to clusters that could be assigned an ICTV taxa. Plots of the cluster composition from the bipartite network analysis were made with ggplot2 (3.1.1) [63] in R with Rstudio. TerL (terminase large subunit) and MCP (major capsid protein) trees were made with hits to TerL VOG families and MCP VOG families encoded by the jumbo phages and reference hits (HMM searches, E-value < 0.001; Supplemental Dataset 2). Reference hits were de-replicated with CD-HIT (-c 0.9) [64] and filtered for size (See Supplementary Methods). Proteins were then aligned with Clustal Omega [65], trimmed with trimAl (-gt 0.1) [66] and constructed with IQ-TREE [67] (TEST model selection with ModelFinder [68]).

### Size fraction and ecological analyses

Metagenomic reads from Tara Oceans were mapped onto the population representatives of the jumbo phage set (535 populations) with coverM (coverm genome; >10% genome covered, min ID 95%), and reads per kilobase per million (RPKM) was then calculated by dividing the number of reads aligned to a phage by the length of the phage in kilobases and then dividing that by the number of reads in the sample in millions, which accounts for differences in phage sequence length and differences in sample sequencing depth. Statistical analyses and plots were carried out in R with vegan (2.5-5) [69], ggplot2, maps (3.3.0) [70], and ggpubr (0.2.4) [71] packages. Community composition was compared between variables using ANOSIMs based on Bray-Curtis distances using both presence/absence and RPKM matrices with a significance p-value cutoff of 0.05. Statistical tests were carried out with the ggplot2 function stat_compare_means(label=“p.signif”).

### Annotation

Amino acid sequences of genes were annotated with HMM searches (Evalue < 0.001) against the Pfam [72] (version 32), eggNOG [73], and VOG databases. Virion structural protein families were identified based on VOG hit descriptions (Supplemental Dataset 3). Consensus annotation was based on Pfam annotations and then the highest bit score between eggNOG and VOG hits. Functions were grouped into larger categories (Supplemental Dataset 3). Clusters from the network analyses were compared for functional composition by averaging the proportion of genes in a functional category using python (version 3) statistics and plotted with ggplot2 in R. Genes with the highest variance between clusters were identified based on the variance in proportion of genomes in a cluster with that gene. Those with over 0.2 variance and a known function were visualized with pheatmap [74] in R.

### Host prediction

Hosts of the jumbo phage bins were predicted based on matching CRISPR spacers, tRNAs, and gene content. CRISPR spacers were predicted on the Genome Taxonomy Database (release 95) [75], metagenome assembled genomes (MAGs) of bacteria and archaea from the metagenomes that the jumbo phage bins derived (provided by Parks et al. 2017 [30]), and on the jumbo phage bins with minCED (derived from reference [76]). Spacers were aligned with BLASTn and matches were >24 bp with <= 1 mismatches [4]. tRNA sequences were predicted with tRNAscan-SE (-bacteria option) [77] on the MAGs and jumbo phage bins. Promiscuous tRNAs [78] were removed (BLASTn hits 100% ID, <= 1 mismatches). Jumbo phage tRNAs were aligned against the MAGs tRNAs and NCBI nr database (BLASTn 100% ID, <= 1 mismatches) [4]. Lastly, hosts were assigned based on the taxonomy of coding sequence matches to the MAGs (BLASTn). Hits to phyla were summed and a putative host phylum had three times the number of hits as the phylum with the next most hits as used in a previous study [4].

## Acknowledgements

We thank members of the Aylward Lab for helpful feedback. This work was performed using compute nodes available at the Virginia Tech Advanced Research and Computing Center.

