## Supplementary material for "Infection strategy and biogeography distinguish cosmopolitan groups of marine jumbo bacteriophages": WeinheimerAylward_marine_jumbo_suppl.pdf

**Supplemental Information**

**Supplemental Dataset 1:** Excel file. Sheet 1: marine jumbo phage genomic features (length, contigs, population representatives), Sheet 2: jumbo bin phage detection results, Sheet 3: host prediction results compiled

**Supplemental Dataset 2:** Excel file. Sheet 1: VOG matrix used to generate network and cluster membership of all sequences in network. Sheet 2: List of VOG families used for MCP, TerL phylogenies.

**Supplemental Dataset 3:** Excel file. Sheet 1: Protein annotation file with hits to EggNOG, VOG,and Pfam; Sheet 2: functional category descriptions; Sheet 3 specifies virion structure VOGs

**Supplemental Dataset 4:** Excel file. Sheet 1: sample metadata; Sheets 2-5 Read mapping results (read counts, fraction covered, RPKM, presence/absence)

**Supplemental Figures**

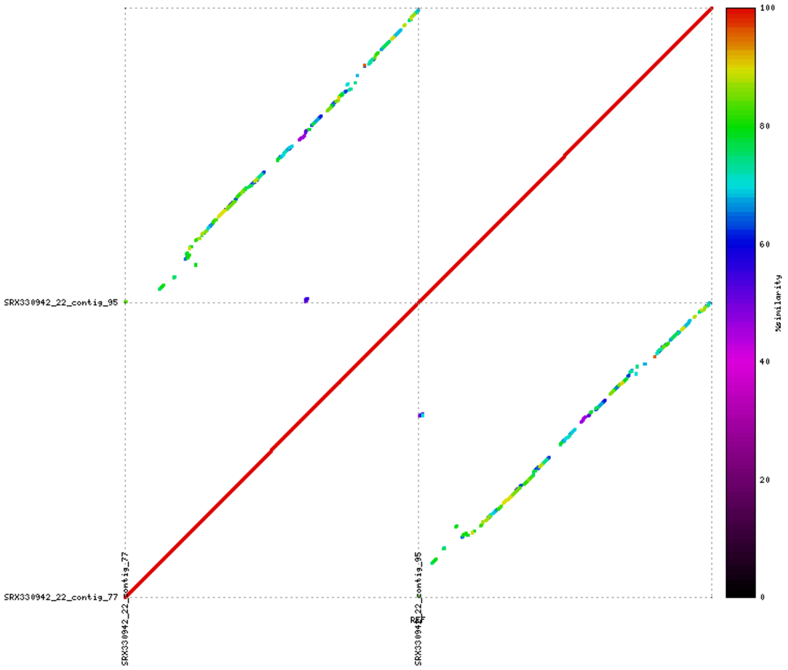


**Supplemental Figure 1.** Example mummerplot of promer alignment between contigs of a single bin. The top right quadrant shows the alignment of the top contig to the bottom contig and the top left quadrant shows the alignment of the top contig to itself. The color of the line corresponds to percent identity. Diagonal lines in the top left and bottom right quadrant show the two contigs of this bin align to each other across their entire lengths with relatively high percent identity, suggesting these contigs belong to two smaller phages, rather than a single jumbo phage genome.

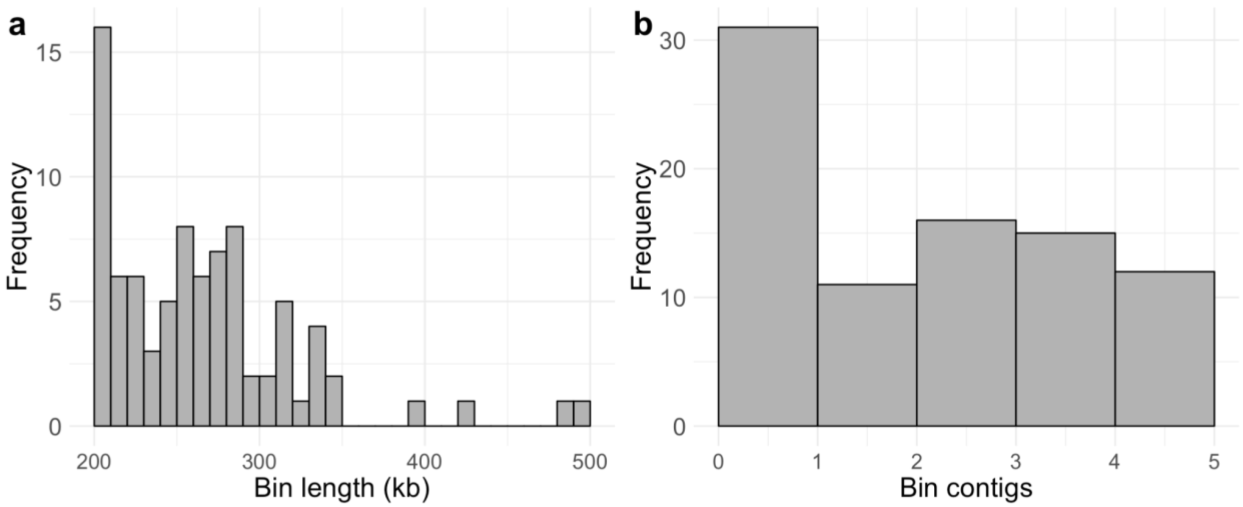


**Supplemental Figure 2. (a)** Histogram of jumbo bin lengths **(b)** Histogram of the number of contigs in each bin

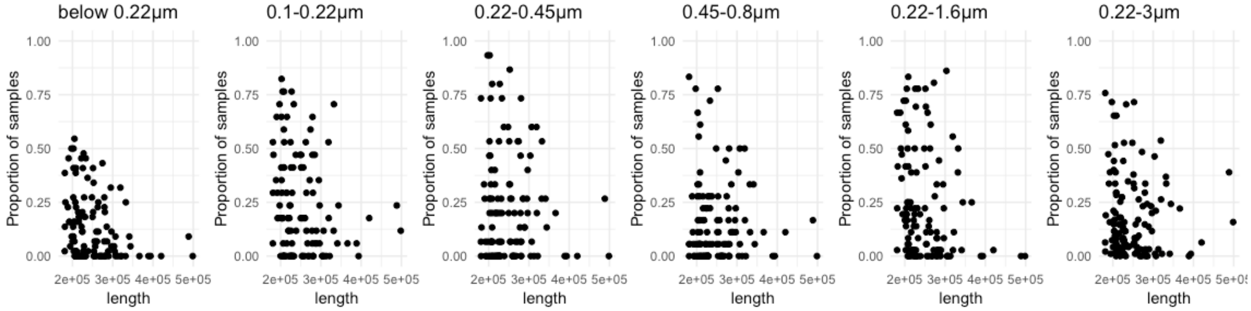


**Supplemental Figure 3.** Scatterplot with proportion of samples at different size fractions that a jumbo phage is present (y-axis) vs. its length (x-axis)

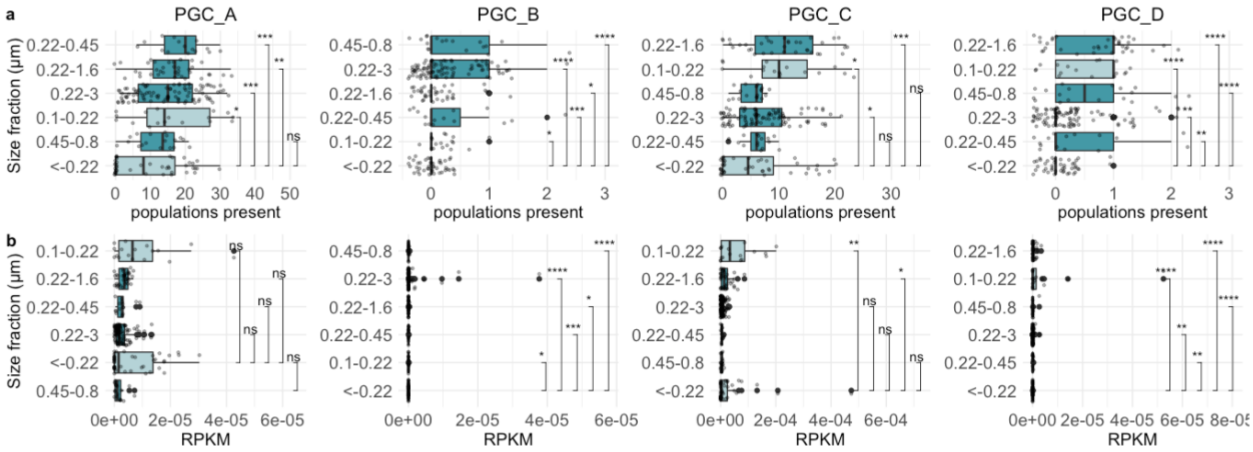


**Supplemental Figure 4. (a)** boxplots for each PGC of the number of jumbo phage populations in each sample of different size fractions sorted by median.**(b)** boxplots for each PGC of the relative abundance of jumbo phages (RPKM) in each sample of different size fractions sorted by median. Significance bars correspond to Wilcox test, with stars corresponding to pvalues < 0.05 and those with pvalues > 0.05 as not significant "ns" (ggplot2 stat_compare_means function).

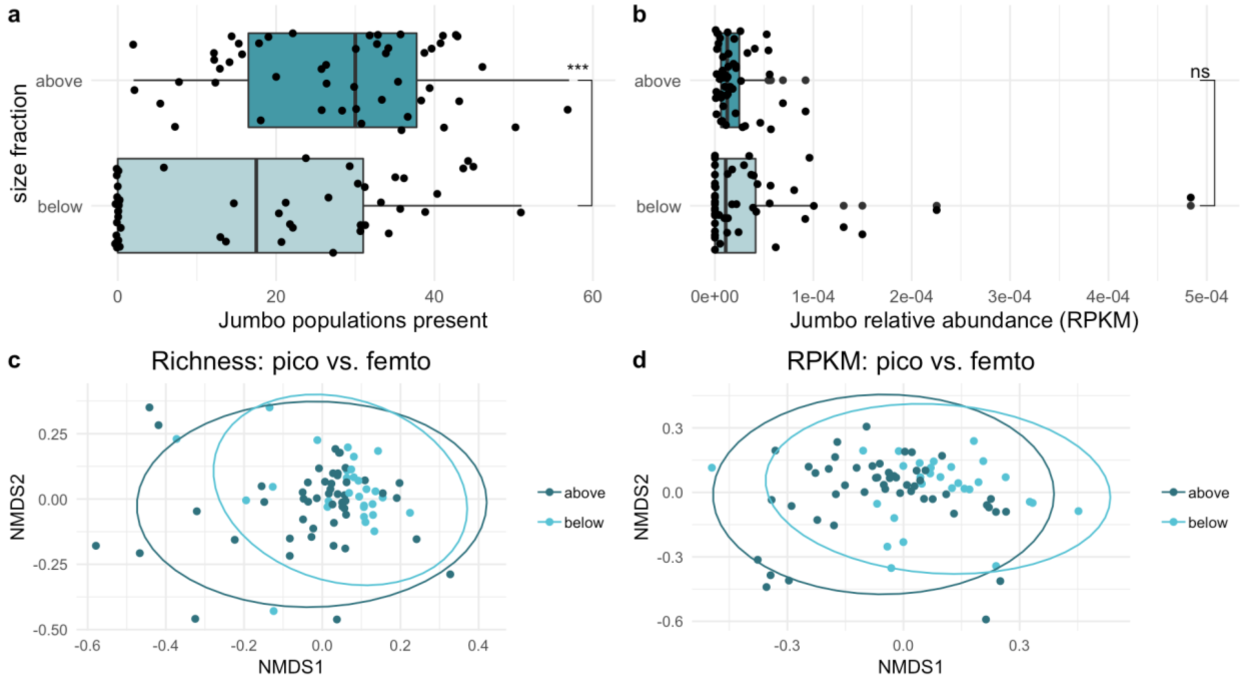


**Supplemental Figure 5.** (**a**) Boxplot of the number of jumbo phage populations present co-collected at the same station and depth but filtered at below 0.22 µm (size fractions "<-0.22µm" or "0.1-0.22µm") and above 0.22 µm (size fraction "0.22-1.6µm" or "0.22-3µm") (**b**) boxplot of the total RPKM of jumbo phages in these samples. Significance bars for a,b correspond to Wilcox test, with stars corresponding to pvalues < 0.05 and those with pvalues > 0.05 as not significant "ns" (ggplot2 stat_compare_means function). (**c**) NMDS plot of samples based on Bray-Curtis distance matrices of jumbo populations' presence/absence (Richness) and communities did not significantly differ between above and below 0.22 (P-value = 0.1229, ANOSIM) (**d**) NMDS plot based on jumbo populations' RPKM; communities significantly differed between above and below 0.22 (P-value = 0.0145, ANOSIM). Ellipses calculated based on multivariate normal distribution.

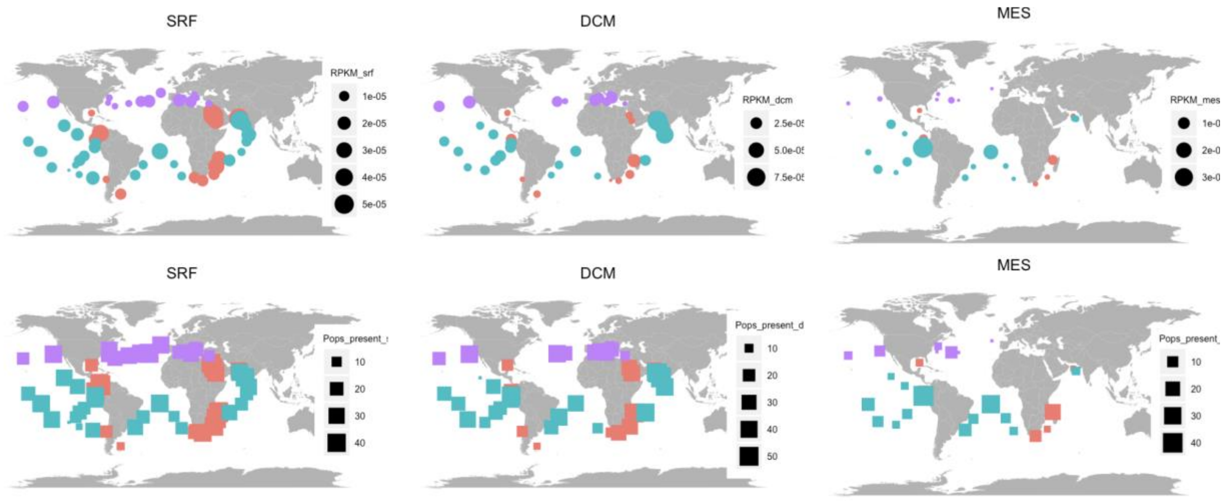


**Supplemental Figure 6.** Distribution of picoplankton fraction at all depths. Upper maps display jumbo abundance (RPKM) in size of dots and bottom maps display jumbo population richness in size of squares. Colors correspond to biome: Coastal - pink, Westerlies - purple, Trades - blue.

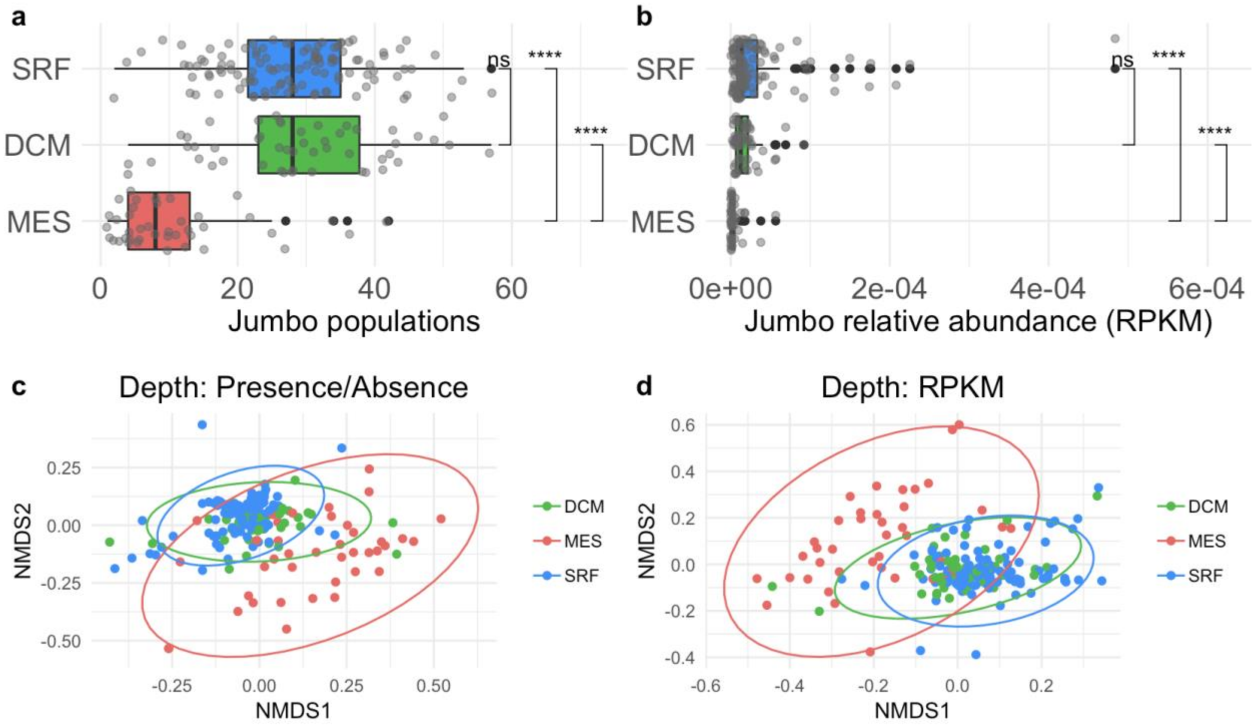


**Supplemental Figure 7. (a,b)** boxplots of jumbo populations present **(a)** and jumbo abundance in RPKM **(b)** in all samples by depth sorted by median abundance. Significance bars for a,b correspond to Wilcox test, with stars corresponding to pvalues < 0.05 and those with pvalues > 0.05 as not significant "ns" (ggplot2 stat_compare_means function). **(c,d)** NMDS plots of jumbo composition in those samples based on Bray-Curtis dissimilarity distances using jumbo populations' presence/absence data **(c)** and jumbo population relative abundance in RPKM **(d)** colored by depth. Green - DCM, red - MES, blue - SRF. Ellipses calculated by multivariate normal distribution. Depths were significantly different using ANOSIM (p-value < 0.05).

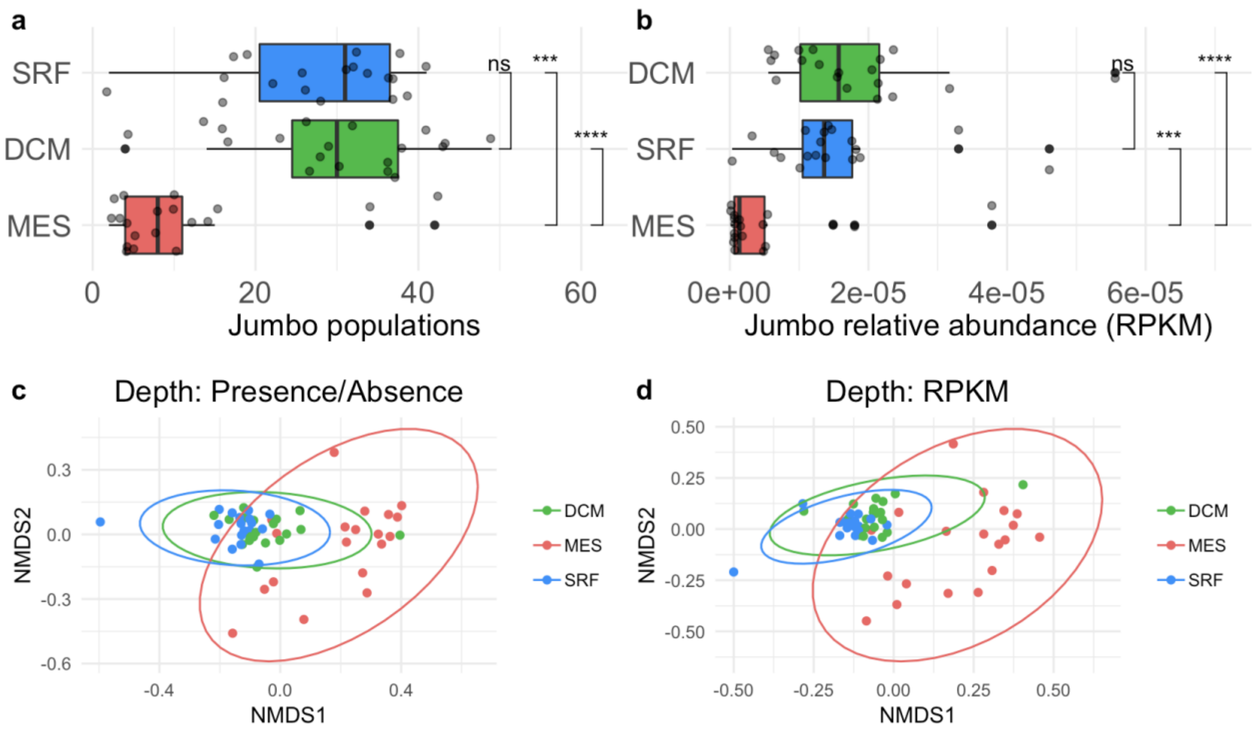


**Supplemental Figure 8. (a,b)** boxplots of jumbo populations present **(a)** and jumbo abundance in RPKM **(b)** in samples of stations co-collected at all three depths in the picoplankton fraction sorted by median abundance. Significance bars for a,b correspond to Wilcox test, with stars corresponding to pvalues < 0.05 and those with pvalues > 0.05 as not significant "ns" (ggplot2 stat_compare_means function). **(c,d)** NMDS plots of jumbo composition in those samples based on Bray-Curtis dissimilarity distances using jumbo populations' presence/absence data **(c)** and jumbo population relative abundance in RPKM **(d)** colored by depth. Green - DCM, red - MES, blue - SRF. Ellipses calculated by multivariate normal distribution. Depths were significantly different using ANOSIM (p-value < 0.05).

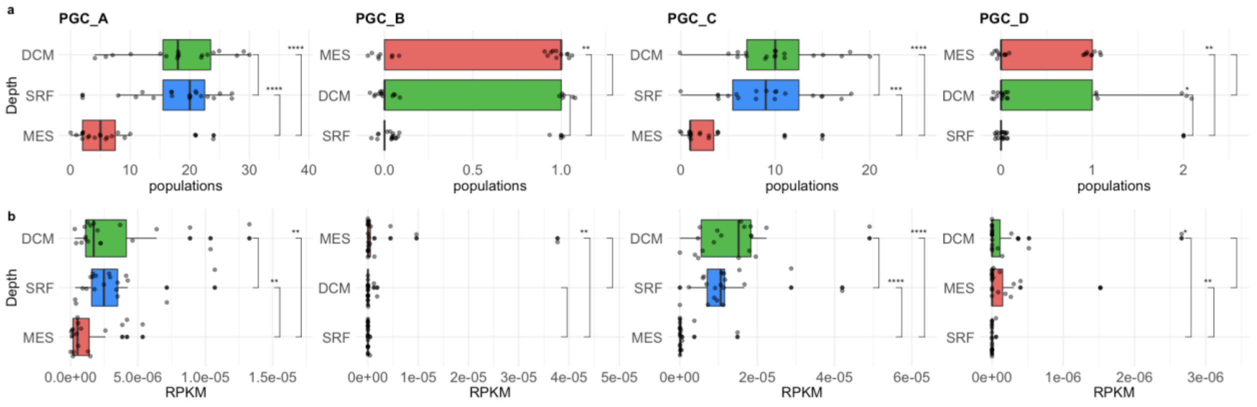


**Supplemental Figure 9. (a,b)** boxplots for PGCs A-D of jumbo populations present **(a)** and jumbo abundance in RPKM **(b)** in samples of stations co-collected at all three depths in the picoplankton fraction sorted by mean abundance. Significance bars correspond to Wilcox test, with stars corresponding to pvalues < 0.05 and those with pvalues > 0.05 as not significant "ns" (ggplot2 stat_compare_means function).

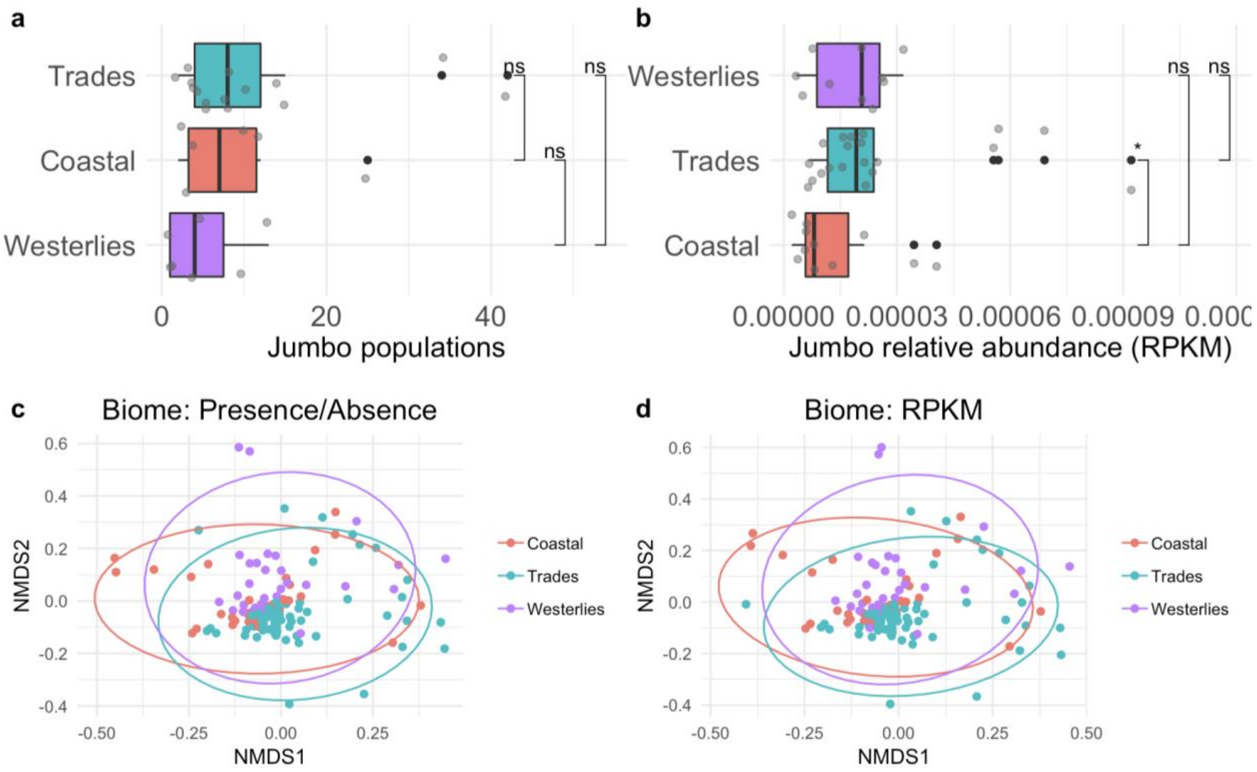


**Supplemental Figure 10. (a,b)** boxplots of jumbo abundance in RPKM **(a)** and jumbo population richness **(b)** in samples co-collected for depth, sorted by median abundance at different biomes. **(c,d)** NMDS plots of jumbo composition in those samples based on jumbo population abundance **(c)** and jumbo populations' presence **(d)** colored by biome. pink - Coastal, blue - Trades, purple - Westerlies. Ellipses calculated by multivariate normal distribution. Biomes were significantly different using ANOSIM (p-value < 0.05, R statistic 0.2). Significance bars in a,b correspond to Wilcox test, with stars corresponding to pvalues < 0.05 and those with pvalues > 0.05 as not significant "ns" (ggplot2 stat_compare_means function).

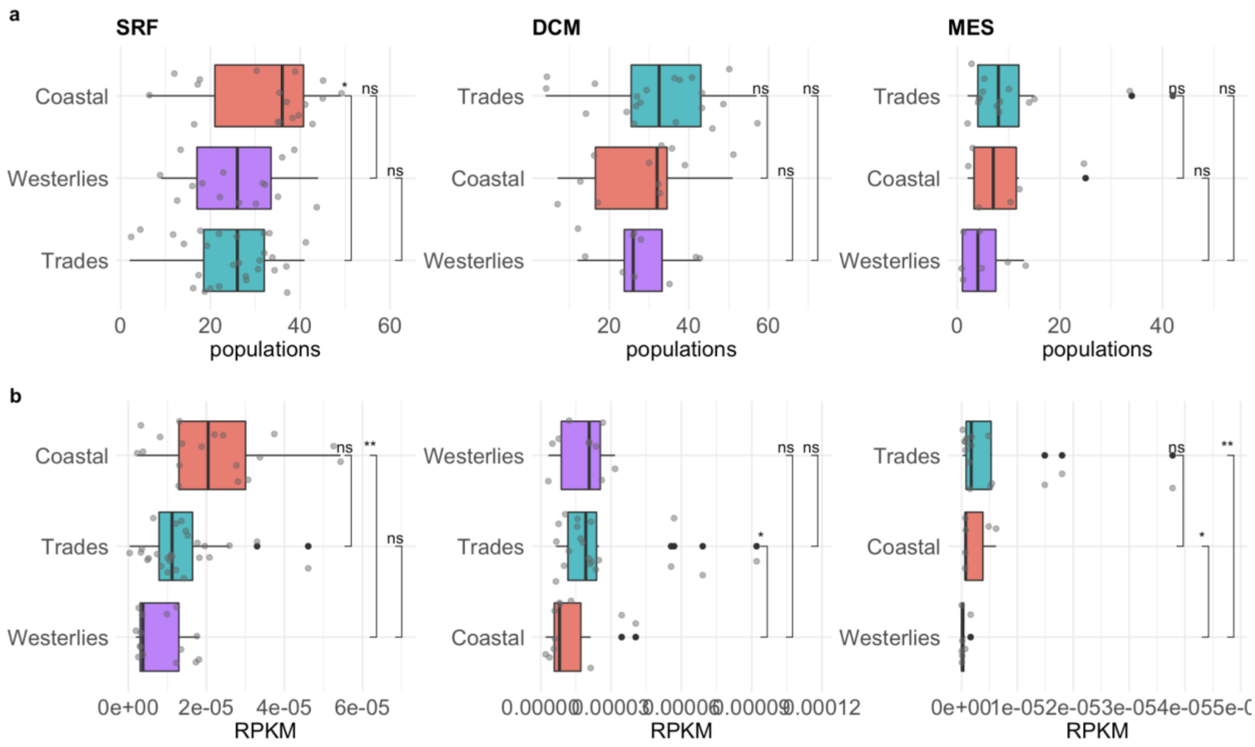


**Supplemental Figure 11. (a,b)** boxplots of jumbo abundance in RPKM **(a)** and jumbo population richness **(b)** in picoplankton samples of each depth separated by biome and sorted by median abundance in the different biomes. Significance bars correspond to Wilcox test, with stars corresponding to pvalues < 0.05 and those with pvalues > 0.05 as not significant "ns" (ggplot2 stat_compare_means function).

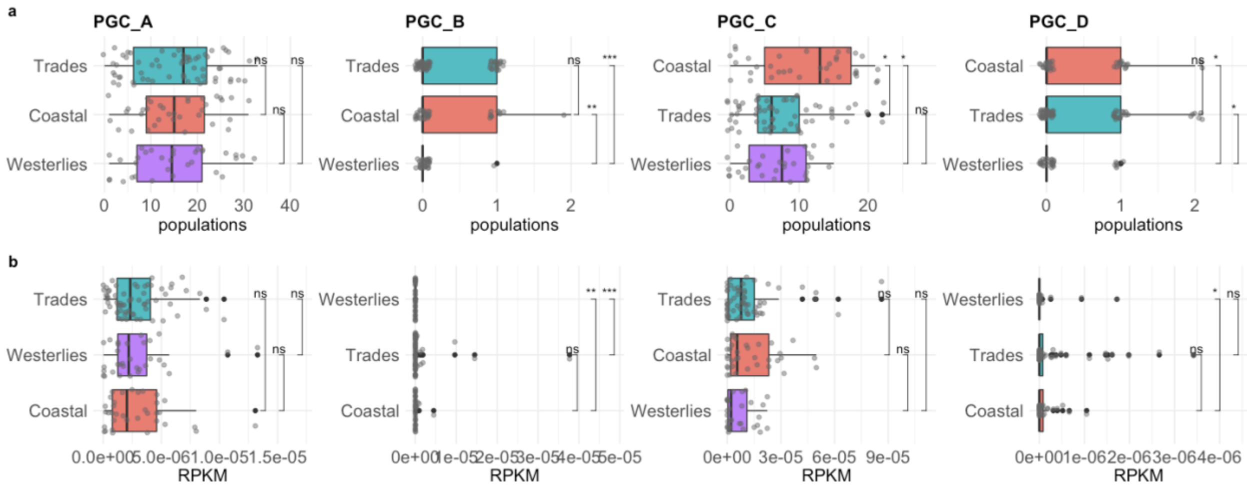


**Supplemental Figure 12. (a,b)** boxplots of jumbo abundance in RPKM **(a)** and jumbo population richness **(b)** in picoplankton samples of each cluster separated by biome and sorted by median abundance in the different biomes. Significance bars correspond to Wilcox test, with stars corresponding to pvalues < 0.05 and those with pvalues > 0.05 as not significant "ns" (ggplot2 stat_compare_means function).

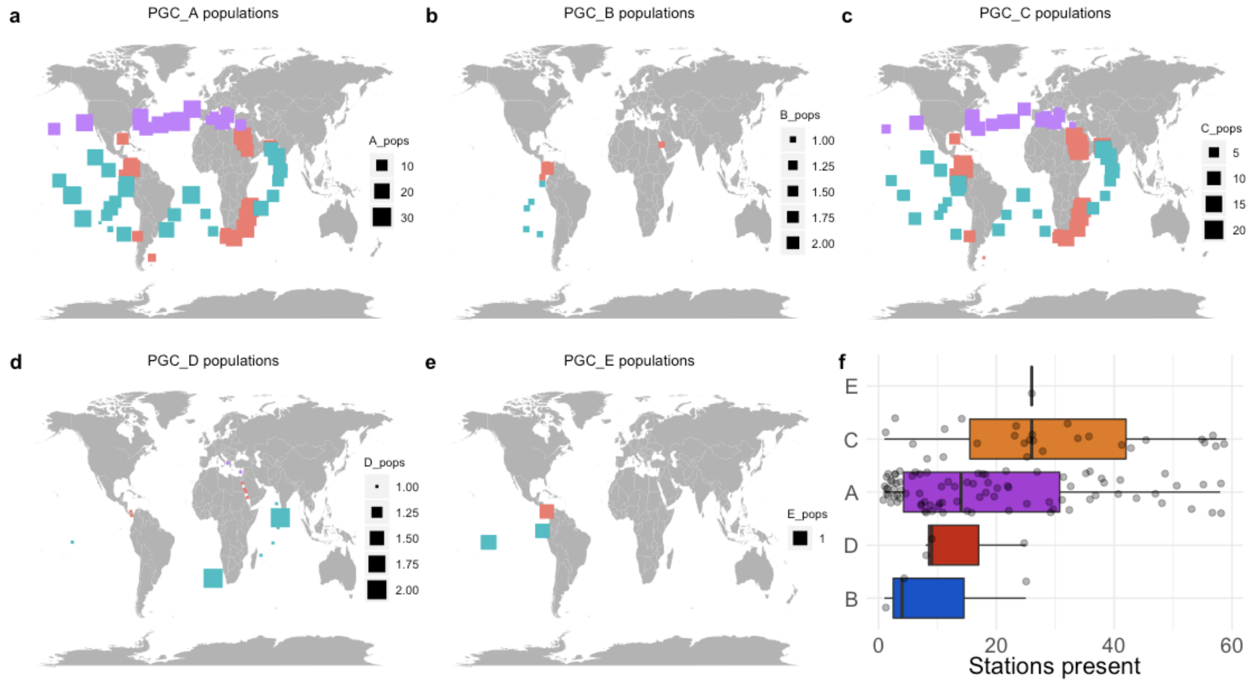


**Supplemental Figure 14.** (a-e) Maps of the number of jumbo populations belonging to the stated PGC in surface samples of the picoplankton fraction, colored by biome. (f) boxplot of the total number of stations (all depths and size fractions) that a jumbo phage was present separated by PGC and sorted by median.

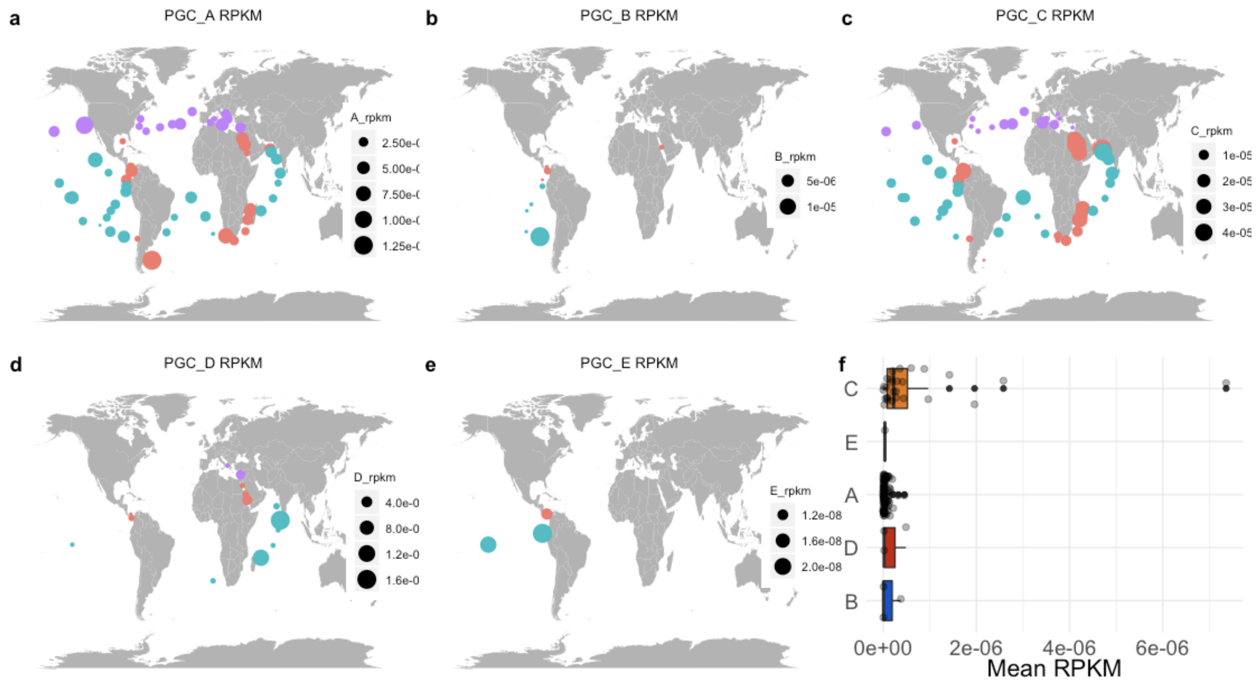

**Supplemental Figure 15.** (a-e) Maps of the relative abundance (RPKM) of jumbo populations belonging to the stated PGC in surface samples of the picoplankton fraction, colored by biome. (f) boxplot of a jumbo population's mean RPKM in all samples (all depths and sizes) separated by PGC and sorted by median.

### Supplemental Methods with References

#### Supplemental Methods

*Binning and screening for non-phage bins.*

Contig sequences and coverage information from 1,545 metagenomes were downloaded from Parks et al 2017 [1]. Contigs were binned with MetaBAT 2 [2] with the options --maxEdge 75 --minS 75 -m 5000, and -s 200000, which resulted in 41,359 bins.

Bins were then filtered for containing a maximum of 5 contigs (1,456 bins remained). We then predicted the proteins on each bin with prodigal [3] using default options. To begin filtering out bins potentially belonging to cells, we removed bins that encoded more than 1 ribosomal protein, which were detected with hidden markov model (HMM) searches via HMMER version 3.2.1) [4] (E value 0.001) against 27 Cluster of Orthologous Groups ribosome protein HMM profiles [5] (1,043 bins remained) (do I put these profiles in Data Availability or Supplemental). Next, we ran a beta version of ViralRecall [6] on the bins to remove bins that had negative scores, which indicate they encode more cellular proteins than viral proteins (673 bins remained).

To address the automated binning complication of strain heterogeneity (cases where contigs binned together based on similar tetranucleotide frequencies and coverage, but actually belong to different viruses), we examined for potential overlapping of conserved regions between contigs by running promer (--maxmatch option) via MUMmer [7], which compares sequences to each other. We then examined this output with mummerplot (--color --png options) for cases where contigs contained extended conserved regions with other contigs in the bin and discarded these bins (example in Supplemental Figure 15).

The remaining 642 bins were then screened for Nucleocytoplasmic Large DNA Viruses (NCLDV) by searching the bin proteins for 8 NCLDV markers with an HMM search (E value 0.001). These eight markers included the following with the minimum bitscore cutoff to be considered a hit in parentheses: A32 (200), D5 (200), SFII (200), mcp (200), mRNAc (200), PolB (500), RNR (200), VLTF3 (200). Additionally, a LASTp [8] was run on the proteins of the 642 bins against RefSeq r99 (E value < 0.001). If the taxonomy of hits to phage proteins outnumbered hits to NCLDV proteins or the number of NCLDV markers was below 2, the bin was considered phage (622 bins remained). Bins were then filtered to remove spurious contigs by removing bins with contigs shorter than 5 kb (610 bins remained). Additionally, we removed bins that contained potentially contaminating contigs based on read mapping coverage (see below).

*Validation of bins with multiple metagenomic read mapping and detection in marine samples*

To further ensure contigs belonging to different phages were not spuriously binned together, we assessed for evenness in contig coverage by mapping reads from different metagenomes to the bins. We used Tara Oceans metagenomes for the mapping [9] so these results could also be used to detect marine jumbo phages. Specifically, we focused on results from samples filtered above 0.22 um to minimize instances of fragmented capsids or free DNA complicating coverage results, which may be more likely in the femtoplankton fraction if the capsid is larger than 0.22 um. Because read mapping evenness can vary in phage genomes due to conserved regions [10], we used mapping results from a reference dataset to benchmark a threshold variation level. For this, we compiled this reference dataset by downloading nucleotide sequences of all complete genomes belonging to the *Caudovirales* order on NCBI's Viral Genomes Portal on July 5, 2020 (referred to as "RefSeq Caudo") and subsetted for jumbo phages ("RefSeq jumbo"); we also included jumbo phage sequences curated by Al-Shayeb et al 2020 [11]. We fragmented these reference jumbo reference sequences with an in-house python script into contigs (1-5) of over 10 kb in length. We then mapped the Tara Oceans metagenomes to this reference set and the bins with multiple contigs (342 bins) with coverM [12] and retained phages with at least 10% covered. Next, we calculated the standard deviation of coverage reported in reads per kilobase per million (RPKM) of the different contigs in a bin with a python script. Reads per kilobase per million is calculated by dividing the number of reads mapped to a sequence by the sequence length in kilobases to account for differences in sequence length between genomes and then dividing that by the million number of reads in the sample to account for differences in read depth between samples. The RPKM tables of the bins and references were split by Tara Oceans depth ("env") type (SRF, DCM, MES). In R (3.5.1) [13] via RStudio (1.1.456). For each depth, we set the maximum standard deviation cutoff to the 95th percentile standard deviation value of the reference RPKM variation (i.e. quantile(reference_srf$std_dev, 0.95). To determine the percentage of samples a bin must have mapped below the reference 0.95 cutoff at each depth, we filtered the bin RPKM table for each depth using percent below the cutoff until the distribution of the standard deviation values for the bins was not significantly different from the reference distribution using a Wilcox test (p-value > 0.05). 310 of the 342 bins passed. 268 bins comprised only one contig, totaling the bins at 578. Based on the read mapping results from samples of all size fractions, a jumbo phage was considered present in a sample if at least 10% of its genome was mapped by the sample. Of the 578 bins that passed the validation test, 107 bins were present in marine samples.

*Validation of bins as phage with phage-detection tools and population clustering with other jumbo phages*

Contigs of the remaining 107 bins were run through VirSorter2 [14] and VIBRANT [15]. For CheckV [16], pseudocontigs were generated using an in-house python script to join the contigs of the bins together with "N"s. First, bins were retained if the VirSorter2 dsDNAphage score of their contigs averaged above 0.9 (75 bins). Next, bins were retained that had a minimum VirSorter2 dsDNA score average above 0.5 and either had been (i) classified as "virus" by VIBRANT or (ii) considered viral by CheckV with genome quality of medium or above. The bins used for subsequent analyses then totaled at 85 bins.

We then compared the bins to other jumbo phages and identified those belonging to the same population, defined by sharing over 80% of genes with at lesat 95% average nucleotide identity. This jumbo phage reference set included those prepared by Al-Shayeb et al. 2020 (336 phages) [11], those available in GenBank compiled by Iyer et al 2021 [17] Cook et al 2021[18] (354 phages), GOV 2.0 (60 phages) [19], ALOHA 2 (8 phages) [20], and one megaphage from the

English Channel [21]. These additional jumbo phage sequences and the phage sequences from RefSeq jumbo are referred to as the "jumbo references", totaling at 852 sequences. Nucleotide and amino acid sequences of genes encoded by the 85 jumbo bins and the 852 jumbo references were predicted with prodigal using the default genome setting for each genome individually (-a,-d options). These genes were then aligned to each other with BLASTn. Bins were considered belonging to the same population if 80% of their genes aligned to another bin's genes with an average nucleotide identity of at least 95% [22]. This analysis resulted in 535 jumbo phage populations, 59 of which contained a jumbo bin generated from this study.

*Bipartite network analysis*

Jumbo bins were clustered with the jumbo references and Caudo RefSeq of all genome sizes based on composition Virus Orthologous Groups (VOG: vogdb.org, downloaded April 14, 2020). Amino acid sequences were searched against HMM profiles in the VOG database via HMM searches ( Evalue < 0.001). A matrix of VOG families as columns and phage as rows was generated from the hmm output with an in-house python script. The matrix was loaded into R and an incidence graph was computed with the R library igraph(1.2.5) [23]. Clusters were then detected with the spinglass algorithm [24] using 50 spins. The spinglass clustering was run 100 times with different seeds. The final clusters were discerned based on the iteration that yielded the highest modularity (seed 544, modularity 0.5856642). Network was visualized with igraph using the Fruchterman-Reingold layout with 5000 iterations (layout.fruchterman.reingold(niter=5000)). Clusters of which the jumbo bins belonged were plotted with ggplot2(3.1.1) [25] in R for composition of RefSeq phages host phyla and dataset origin; figures were joined with ggpubr(0.2.4) and Inkscape(v 0.92).

*MCP and TerL Phylogenies*

All major capsid protein (MCP) and terminase large subunit HMM profiles were compiled from vogdb.org (release 98). Proteins of the jumbo bins, jumbo references, and RefSeq Caudo were searched against these databases with HMM searches (-E 0.001 flag). To reduce the reference dataset to facilitate phylogenetic analyses, we first took the best hit of MCP and TerL encoded by a reference based on bitscore and then clustered the reference hits with cd-hit[26] using a 90% ID cutoff (-c 0.9 option). Lastly, to improve alignment quality, proteins from both the references and jumbo bins from this study were removed that were less than two standard deviations below the median length encoded by the references, which was 96 amino acids (aa) for the MCP and 170 aa for TerL. For the bins, only the top hit of these remaining MCP and TerL were included in the phylogenetic analysis, resulting in 74 MCP bin hits and 80 TerL bin hits.

In total, 1,195 MCP protein sequences and 1,367 TerL sequences were aligned separately with Clustal Omega [27]. The alignments were then trimmed using trimAl (parameter -gt 0.1) [28]. A tree was reconstructed with this alignment using IQ-TREE [29] with ModelFinder [30] to select the best fit model according to the Bayesian Information Criterion, which was VT+F+G4 for the MCP alignment and Blosum62+F+G4 for the TerL alignment and 1,000 ultrafast bootstrap replicates. Trees were visualized in iTOL (v5) [31] and jumbo bins were colored with network cluster. Figures were joined with ggpubr and Inkscape.

*Annotation*

Amino acid sequences of jumbo phages were searched with HMM searches against HMM profiles of the EggNOG 5.0[32] (Evalue <0.001 ), VOG (release 98), and Pfam (Pfam-A, version 32) [33] databases. To identify virion structural proteins, structural proteins in VOG were manually identified (Supplemental Dataset 2). A consensus annotation of a protein was determined based first by Pfam hit because these functions are well-curated [33], then by VOG or EggNOG hit based on bitscore. Pfam does not assign functional categories, so a gene's functional category was based on the category of its EggNOG hit or virion structure designation. These categories were subsequently merged into broader categories (Supplementary Dataset 2). Functions that had multiple EggNOG categories (i.e. NK) were tallied individually. Stacked barplots of the functional composition of each jumbo phage cluster were based on the average proportion of genes belonging to the category and plotted in R with ggplot2. Genes with known functions that drove variation between clusters A-D of this study were identified by first calculating the proportion of genomes which encoded a given gene in each cluster and then calculating the variance of the proportion of genomes among the clusters in R. Genes with variance above 0.2 and known functions were retained for heatmap visualization made with pheatmap in R.

Group 1.0 jumbo phages, belonging to cluster B in this study, are known to encode a divergent family B DNA polymerase. As this gene has not been included in the databases examined, we identified the HMM profile of the VOG family corresponding to this divergent family B DNA polymerase by searching a reference sequence for this gene (YP_009153312.1) with an HMM search (-E 0.001) against the VOG database, which was VOG09941 (bitscore > 1000). We then compared the bitscore of genes that hit to this VOG with the classic family B DNA polymerase (PF00136) to identify the occurrence of the divergent family B DNA polymerase in the phages.

*Distribution analyses*

To examine the distribution of populations of jumbo phages in the ocean, we mapped reads from the Tara Oceans metagenomes used in the bin validation, but excluded Polar samples as there were only 5 available in this set. Reads were mapped onto the representative sequences of the 535 jumbo phage populations that were selected based on highest length with coverM (coverm genomes; >10% genome covered; min ID 95%). To compare mapping results between phages and samples, reads per kilobase per million (RPKM) was then calculated by dividing the number of reads aligned to a phage by the length of the phage in kilobases and then dividing that by the number of reads in the sample in millions, which accounts for differences in phage sequence length and difference in sample sequencing depth. Statistical tests and plots of the mapping results were carried out in R with the stat_compare_means(label="p.signif") function in ggplot2 to compare samples between biomes, fractions, and depth in richness and abundance of jumbo phage populations. Compositional differences of jumbo phage between samples based on both presence/abundance and RPKM matrices were compared with ANOSIMs in R using the anosim function from the package vegan(2.5-5) [34] (distance="bray",permutations=9999). Maps were plotted in R with the maps [35] and ggplot2 libraries. Boxplots were plotted in R with ggplot2 and plyr(1.8.4) [36] to order axes. Non-metric dimensional scaling plots were generated in R based on Bray Curtis dissimilarity matrices (vegdist (method="bray)) using the metaMDS vegan function and visualized with ggplot2; ellipses were calculated with stat_ellipse(type="norm").Figures were joined with ggpubr in R.

*Host prediction*

Hosts were estimated for the bins based on CRISPR spacers, tRNAs, and the taxonomy of genes. CRISPR spacers were predicted for the Genome Taxonomy Database (release 95)[37] and metagenome assembled genomes (MAGs) of bacteria and archaea from the metagenomes of which the jumbo phages derived by Parks et al 2017 [1], as well of the jumbo bins. All spacers were aligned to the jumbo bins and hits were at least 24 basepairs in length with <= 1 mismatch [11]. No hosts could be assigned with this approach, and no jumbo bins targeted other jumbo bins. tRNAs were predicted on the jumbo bins and the same MAGs set with tRNAscan-SE [38] (-bacteria option). Promiscuous tRNAs were downloaded from Paez-Espino et al. 2016[39] and removed based on BLASTn hits (100% ID, <= 1 mismatches). Jumbo tRNAs were then aligned against the MAGs tRNAs with BLASTn and matches were considered with 100% ID and no more than one mismatch. Jumbo phage tRNA sequences were also searched against NCBI's nonredundant database using the BLASTn webserver and matches were retained with the same criteria. Finally, hosts were assigned based on the taxonomy BLASTn hits to the coding sequences of the MAGs. A putative host phylum was considered if a phylum had three times as many hits than the phylum with the next most hits [11].

*References*

11. Al-Shayeb B, Sachdeva R, Chen L-X, Ward F, Munk P, Devoto A, et al. Clades of huge phages from across Earth's ecosystems. *Nature* 2020; **578**: 425–431.
12. wwood. GitHub - wwood/CoverM: Read coverage calculator for metagenomics. https://github.com/wwood/CoverM. Accessed 23 Jul 2021.
13. R Core Team. R: A Language and Environment for Statistical Computing. 2019. R Foundation for Statistical Computing, Vienna, Austria.
14. Guo J, Bolduc B, Zayed AA, Varsani A, Dominguez-Huerta G, Delmont TO, et al. VirSorter2: a multi-classifier, expert-guided approach to detect diverse DNA and RNA viruses. *Microbiome* 2021; **9**: 37.
15. Kieft K, Zhou Z, Anantharaman K. VIBRANT: automated recovery, annotation and curation of microbial viruses, and evaluation of viral community function from genomic sequences. *Microbiome* 2020; **8**: 90.
16. Nayfach S, Camargo AP, Schulz F, Eloe-Fadrosh E, Roux S, Kyrpides NC. CheckV assesses the quality and completeness of metagenome-assembled viral genomes. *Nat Biotechnol* 2021; **39**: 578–585.
17. Iyer LM, Anantharaman V, Krishnan A, Maxwell Burroughs A, Aravind L. Jumbo Phages: A Comparative Genomic Overview of Core Functions and Adaptions for Biological Conflicts. *Viruses* . 2021. , **13**: 63
18. Cook R, Brown N, Redgwell T, Rihtman B, Barnes M, Stekel DJ, et al. INfrastructure for a PHAge REference Database: Identification of large-scale biases in the current collection of phage genomes.
19. Gregory AC, Zayed AA, Conceição-Neto N, Temperton B, Bolduc B, Alberti A, et al. Marine DNA Viral Macro- and Microdiversity from Pole to Pole. *Cell* 2019; **177**: 1109–1123.e14.
20. Luo E, Eppley JM, Romano AE, Mende DR, DeLong EF. Double-stranded DNA virioplankton dynamics and reproductive strategies in the oligotrophic open ocean water column. *The ISME Journal* . 2020. , **14**: 1304–1315
21. Michniewski S, Rihtman B, Cook R, Jones MA, Wilson WH, Scanlan DJ, et al. Identification of a new family of 'megaphages' that are abundant in the marine environment. *bioRxiv* . 2021. , 2021.07.26.453748
22. Brum JR, Ignacio-Espinoza JC, Roux S, Doulcier G, Acinas SG, Alberti A, et al. Ocean plankton. Patterns and ecological drivers of ocean viral communities. *Science* 2015; **348**: 1261498.
23. igraph – Network analysis software. http://igraph.org. Accessed 23 Jul 2021.
24. Reichardt J, Bornholdt S. Statistical mechanics of community detection. *Phys Rev E Stat Nonlin Soft Matter Phys* 2006; **74**: 016110.
25. Wickham H. ggplot2. *Wiley Interdisciplinary Reviews: Computational Statistics* . 2011. , **3**: 180–185
26. Fu L, Niu B, Zhu Z, Wu S, Li W. CD-HIT: accelerated for clustering the next-generation sequencing data. *Bioinformatics* 2012; **28**: 3150–3152.
27. Sievers F, Wilm A, Dineen D, Gibson TJ, Karplus K, Li W, et al. Fast, scalable generation of high-quality protein multiple sequence alignments using Clustal Omega. *Mol Syst Biol* 2011; **7**: 539.
28. Capella-Gutiérrez S, Silla-Martínez JM, Gabaldón T. trimAl: a tool for automated alignment trimming in large-scale phylogenetic analyses. *Bioinformatics* 2009; **25**: 1972–1973.
29. Nguyen L-T, Schmidt HA, von Haeseler A, Minh BQ. IQ-TREE: a fast and effective stochastic algorithm for estimating maximum-likelihood phylogenies. *Mol Biol Evol* 2015; **32**: 268–274.
30. Kalyaanamoorthy S, Minh BQ, Wong TKF, von Haeseler A, Jermiin LS. ModelFinder: fast model selection for accurate phylogenetic estimates. *Nat Methods* 2017; **14**: 587–589.
31. Letunic I, Bork P. Interactive Tree Of Life (iTOL) v5: an online tool for phylogenetic tree display and annotation. *Nucleic Acids Res* 2021; **49**: W293–W296.
